# Migration routes and wintering sites of the Aquatic Warblers *Acrocephalus paludicola* breeding in Lithuania and North Belarus

**DOI:** 10.1101/2023.10.10.561376

**Authors:** Martin Flade, Simeon Lisovski, Vytauras Eigirdas, Benedikt Giessing, Fréderic Jiguet, Oskars Keišs, Maxim Nemtchinov

## Abstract

For the effective protection of the globally threatened Aquatic Warbler in its entire range, knowledge of the migration and wintering areas as well as their current conservation status is of great importance. The question on where the different breeding population overwinter where (connectivity) equally important. This is also essential when new breeding populations are re-established in restored breeding sites through translocation from other, distant areas, as recently performed in Lithuania. From previous geolocator studies in central Ukraine and south-western Belarus, as well as from ringing data, it was known that these populations overwinter in Mali (Inner Niger Delta IND) and southern Mauritania. In addition, at least some birds of the NE-Polish breeding population overwinter in the Senegal delta (ring recovery). In 2018 and 2019, the migratory routes, stopover and wintering sites of two breeding populations at the north-western distribution range border in N-Belarus (29 males) and Lithuania (31 males) were investigated using light-level geolocators. All 19 recovered data loggers recorded almost complete annual cycles from mid-July to at least early April. Migration and overwintering sites of both populations did not differ significantly. Most (16 out of 19) birds also spent the winter in the Inner Niger Delta and surrounding wetlands within and around Mali. Only one bird from Belarus hiberbated in the Senegal Delta. One of the Lithuanian birds overwintered in northern Burkina Faso, another northern Nigeria. In addition, important details about the course of migration routes, timing and resting areas could be obtained. For example, the outstanding importance of Morocco and northern Algeria as resting areas on spring migration became obvious. The investigations applying geolocators resulted in a very complex and differentiated picture of the migration and stopover pattern of adult male Aquatic Warblers.

## 1. Introduction, framework and aim of the study

The Aquatic Warbler (AW) is an endemic European breeding bird and one of the rarest passerine species of Europe. During the breeding season it inhabits meso-to eutrophic, shallowly flooded large open sedge fen mires, *Cladium* marshes, coastal brackish marshes with low reeds, and similarly structured wet sedge meadows in floodplains used for hay-making (Tanneberger et al. 2018). During migration through West and Southwest Europe AW uses mostly coastal wetlands at river estuaries and lagoons as stopover habitats (Le Nevé at al. 2018) and is wintering in wetlands of the western Sahel south of the Sahara (Tegetmeyer et al. 2018).

Due to large-scale habitat degradation of >95 % of the former breeding sites through drainage, agricultural use and abandonment, the AW is now one out of three passerine breeding bird species of continental Europe which are threatened at a global level, classified as vulnerable (Birdlife International 2015). The remaining global population is estimated at 11,000 singing males (Flade et al. 2018), which are currently concentrated at less than 50 breeding sites in four countries only (Belarus, Ukraine, Poland, and Lithuania). Some 70-78 % of the population is concentrated in just four key breeding sites (Flade et al. 2018). In order to secure the global population, the whole annual cycle of the species has to be considered. The knowledge about connectivity between specific breeding populations and wintering sites, and knowledge about the importance of specific stopover sites on autumn and spring migration is of very high importance.

Aim of this study was to identify the migration routes and wintering sites of the Lithuanian and North-Belarusian AW populations. Since the discovery of wintering AW in Djoudj, Senegal delta (Flade et al. 2011), follow-up studies on the ground in Senegal, Mauritania, Mali and Gambia, studies by satellite image analyses and on stable isotopes in AW feathers, have most likely identified the main African wintering grounds (Flade & Lachmann 2008, Flade et al. 2011, Buchanan et al. 2011, Oppel et al. 2011, Poluda et al. 2012, Salewski et al. 2013, 2019). They are located between the Senegal delta in the west, the Inner Niger Delta in Mali in the east and southwards reaching possibly the northernmost parts of Ivory Coast and Ghana (Flade et al. 2011, Salewski et al 2018).

The next step was to investigate the connectivity between breeding populations and wintering sites. Results of two former studies with geolocators on AW populations in central Ukraine (Supii) and Southwest Belarus (Dikoe) showed that these birds spend the winter in western and central Mali (Salewski et al. 2013, 2019). A recovery of a male AW, ringed in February 2011 in the Inner Niger Delta (Mali), as a breeding bird in Supii (central Ukraine) in the same year confirmed this result (Poluda et al. 2012).

It was assumed that the AW wintering in Djoudj/Senegal delta belong to the northwestern-most populations in Poland and probably Lithuania. This hypothesis is supported, yet not confirmed, by a ring recovery of a female AW wintering in Djoudj as breeding bird in the Biebrza marshes (Poluda et al. 2012).

The question, whether Lithuanian AW spend the winter in different wintering sites than AW of S-Belarus and Ukraine is of uttermost importance with respect to the successfully piloted translocation of birds from S-Belarus to Lithuania within the running EU-LIFE Project MagniDucatusAcrola (Morkvenas et al. in prep.). If this was the case, birds from a population wintering in Mali would have been translocated into a range where probably AW spend the winter in Senegal.

The knowledge on connectivity between breeding and wintering sites is also important with respect to the effects of differing conditions during wintering on the breeding population dynamics.

## 2. Methods of field work and data analysis

With a body mass of c. 11-12 g, the Aquatic Warbler cannot carry a satellite tag to accurately track its migration routes. Recently, migration flyways of the species were studied using durable and lightweight (0.6 g) light-level geolocators.

The first two geolocator studies on AW in Ukraine and Belarus have proven that the method can lead to useful results (Salewski et al. 2013, 2019). The return rate of birds was not different to the control groups and the birds have been in a good physical condition at recapture. In both studies it was possible to recapture 20-25 % of the tagged birds. More severe problems were of technical nature: limited precision of localisations and malfunction of geolocators. Most geolocators stopped with data storage too early, mostly in autumn, and only two devices measured the complete annual cycle. For our study we therefore used newly developed geolocators from Migration Technology Ltd. (UK), which are even smaller and lighter (0.36-0.38 g) than the formerly used devices (0.6 g) and are known to deliver a higher precision of localisations due to a light sensor recording a higher range of light intensities, and due to less malfunctions.

Only adult males of AW were fitted with geolocators due to their highest probability of recovery (same approach as in Salewski et al. 2013, 2019). Males likely have a higher survival rate than juvenile birds and are much easier to detect and to catch in the following year, when they are actively singing in the breeding grounds.

Based on the past experiences, we used 60 geolocators, 29 in N-Belarus and 31 in Lithuania. Adult males were attracted using tape lures and captured with mist nets. This was done in late June to early July 2018 towards the end of the active song period of the males, in order to minimise disturbances to mating and breeding of the birds. In May 2019 the birds were recaptured (one bird even one year later in June 2020). Singing males at the breeding sites were inspected with a telescope and all ringed males, potentially wearing a geolocator, were recaptured with mist-nets. Trapping of AW was exclusively focussed on potential geolocator birds in order to minimise disturbance.

According to the manufacturer information, the average durance of data logging was 10 months. In order to save power and to cover both migration periods, all devices were programmed to start data logging only on 16^th^ July, 2018.

Harness loop sizes of geolocators differed within a limited range (in our case, width/span of stretched loops has been 33.5-38.5 mm). In order to adjust the harness loop sizes as accurate as possible to the constitution of the respective individual, we measured the loop sizes of all 60 geolocators with callipers and assigned them to size classes (S = small, M = medium, L = large). Thus, we were able to deploy the best fitting geolocator to each captured AW by means of the parameters wing length, weight and ‘feeling in the hand’ of the birds. This additional effort likely was worthwhile, since quite different to the former studies (Salewski et al. 2013, 2019) we found no indication that any AW lost its geolocator or suffered any skin injury by the harness. We did neither find any bird that lost its geolocator during the first days, nor found any ringed AW in May 2019 that had lost its geolocator in the meantime. Considerably improved selection of harness material (‘stretch magic’) could be an additional reason for this successful campaign.

Light-intensity data were recorded at 5 minute intervals and analyzed using a threshold method (Lisovski et al., 2020). Sunrise and sunset events (twilight events) were identified on log transformed light data and a threshold of 0.5 log lux, using the R package TwGeos (Lisovski, Sumner, & Wotherspoon, 2015). Twilight events recorded at the known deployment site during periods after deployment and before recapture where used for calibration, i.e. to estimate the error distribution of twilight events and the individual reference sun elevation angle (position of the sun when twilight events where detected). Migratory movements were identified as sudden and directed changes of consecutive sunrise or sunset events (for details on the methods see Lisovski et al., 2020). Periods of residency with a minimum of three consecutive twilight events were then identified as periods surrounded by migratory movements. For the final track estimation, we used a Bayesian approach from the R Package *SGAT* (Wotherspoon, Sumner, & Lisovski, 2013) allowing to incorporate the twilight events, their error distribution (gamma density distribution), the information on periods of movement and residency, a spatial probability mask, and an expected flight speed distribution (gamma distribution with shape = 1 and rate = 0.05). Altogether and via Markov chain Monte Carlo (MCMC) simulations, the method aims to refine tracks, providing most likely paths with credibility intervals. The *groupedThresholdModel*, providing a single location estimate for periods of residency, was chosen. For periods of defined movements, no mask to spatially restrict location estimates. For periods of residency, location estimates were spatially directed according to a relative likelihood with a probability of 1 for inland locations, −1000 for locations at sea. During the MCMC simulation, the first and last site of residency (in case the logger was still recording light on return) were fixed to the deployment site. We first ran a *modifiedGamma* model (relaxed assumptions) for 1000 iterations to initiate the model, before tuning the model with final assumptions/priors with three runs each containing 300 iterations. Finally, the model was run for 2000 iterations to ensure convergence. Median location estimates and 95% CrI were calculated using the entire final MCMC chain (i.e. each location estimate was based on 2000 estimates from within the MCMC chains).

Based on the analysis of stationary periods, we extracted the departure and arrival dates at the deployment site. Stopover periods were identified prior to the MCMC simulation and based on changes in the sunrise and sunset times. The *groupedThresholdModel* in SGAT estimates one location for each stationary period, based on all twilight times within that period, and one location for each twilight during movement. Locations from the most likely track were used for plotting and further analyses. It has to be noted that the accuracy of localisations solely based on the information obtained from geolocators is quite limited. Moreover, the precision is lowest around equinox (22^nd^/23^rd^ September, 20^th^ March). Assumed stopover sites indicated in Annex 1-4 have therefore be treated as ‘informed guesses’. We attempted to match geolocator locations from stopovers of >2.5 days to specific known AW stopover sites according to ringing and ring recovery data as well as confirmed field observations, or, if confirmed sites are absent, to assumed larger patches of suitable habitat like coastal lagoons, river estuaries, inner river deltas and lakes in the Sahara Desert, wetlands with grassy vegetation or reedbeds (for details of assignments see annex 1-4). Information on the occurrence of such sites has been obtained in Europe from knowledge of national and regional experts (members of the Birdlife International Aquatic Warbler Conservation Team AWCT) and in Africa from AWCT expeditions to Senegal and Mauritania in 2007 and 2008 (Flade et al. 2011), from expeditions of the French group ACROLA to Mauritania and Mali in 2012 (Foucher et al. 2013), and by means of public available Google satellite images. We assessed the probability of correct assignment (high – medium – low – no; see annex 1-4) according to the following criteria: Distance of geolocator localisation to the next known stopover site or the next larger area of probably suitable habitat, confidence interval of the geolocator coordinates, and number of alternative options (possible sites). If there was only one known stopover site at a distance of less than 30 km, probability was always assessed es high, in a distance of up to 80 km (exceptionally up to 120 km) mostly as medium and at larger distances up to c. 150 km as low. For potential patches of suitable habitat without confirmed AW records, or if there occurred several alternative options, or if the confidence interval of geolocator coordinates was very wide, the probability of correct assignment was assessed to be lower.

The state of knowledge about the AW as of 2018 is comprehensively summarised and discussed in the Aquatic Warbler Conservation Handbook (Tanneberger & Kubacka 2018). When discussing the results of our recent study, we therefore refer mostly to the respective chapters of the handbook as basis.

## 3. Study Sites

### 3.1 Selection of study sites

Since AW fitted with geolocators must be recovered and recaptured after one year, the studied populations should be sufficiently small and, on the other hand, sufficiently large (ideally 50-150 singing males), and should be stable and geographically isolated, so that the chance of recovery of the birds is reasonably high. Therefore, the sites Tyrai and Alka polder at and close nearby the Curonian Lagoon in Lithuania, representing the northernmost known AW breeding sites on the world, and Servech in the north of Belarus near the Lithuanian border have been chosen as study sites. Each of these sites held between 30 and 80 singing males in the years previous to our study. The distance from Tyrai and Alka polder (and the neighboring small population at the Nemunas delta) to the next bigger AW population of >10 singing males was 210 km (Biebrza marshes in Poland); the distance from Servech to the next bigger AW populations was 330 km (Biebrza marshes in Poland, Dzikoe and Sporovski in SW-Belarus) (Flade et al. 2018).

**Servech**: Mire at the western and north-eastern bank of Lake Servech, S of Dzyerkawshchyna, SW Glubokae, Vicebsk Region, Belarus; Lake: 55.0338 N, 27.5691 E

**Tyrai**: NNE Dreverna between Vilhelmo Kanalas and the Curonian Lagoon, Lithuania; 55.5318 N, 21.2201 E

**Alka polder**: W Alka, NNW Silute, Lithuania; 55.4327 N, 21.3808 E

### 3.2 Effects of varying habitat conditions in the study sites

To assess the habitat suitability of the study sites during the breeding season of the two study years, some information on the year-specific situation is given. This is important with respect to the local phenology of AW in these years (song period/singing activity and timing of broods), which is affecting capturing probability of males as well as recovery rates in the next year.

In early May 2018, the water table in the Servech mire (Belarus) was still very high, >40 cm above ground. Probably this caused a delay in the beginning of the first broods. During our visit in late June the situation had changed completely: there was not any water above the ground. Between 26^th^-28^th^ June there were still many females uttering alarm calls (at least 8 seen), indicating that still nestlings were fed. In one nest found on 26^th^ June, the nestlings were c. 6 days old. That means that beginning of nest construction was in late May/early June, start of incubation on c. 6^th^ June.

The singing activity of males dropped dramatically on 27^th^ June. On 28^th^ June there was only extremely low activity, and males were much more active in the morning than in the evening. Amazingly, despite normally almost all males are singing during sunset (Glapan 2018), it was completely silent during sunset of 28^th^ June. Our interpretation is, that the males perceived, that most females were still occupied with raising nestlings and thus were not ready for mating, and that the season was already rather late or too late to initiate a second clutch. For this reason, we decided not to spend too much time for catching a 30^th^ geolocator bird in Belarus and to use the remaining 31 geolocators in Lithuania.

As 2019 was the second very dry year after 2018, water table of the lake and in the adjacent mires was low in spring. There was no open water on the ground of the sedge fen. Despite that, in the small mire the abundance of AW and Common Snipe *Gallinago gallinago* seemed to be nearly at average. On the other hand, densities of other typical fen mire birds like Sedge Warbler *Acrocephalus schoenobaenus*, Reed Bunting *Emberiza schoeniclus* and especially Meadow Pipit *Anthus pratensis* were very low.

Spotted Crake *Porzana porzana* and Water Rail *Rallus aquaticus* were almost absent. The big mire was completely without surface water too, and abundances of AW and Sedge Warbler were low. The Meadow Pipit was completely absent. Altogether, the habitat conditions in Servech seemed to be suboptimal for AW and its abundance lower than average.

In contrast to Servech, the Tyrai mire in Lithuania is less dependent on precipitation, because its water table is mainly influenced by the wind tide of the Baltic Sea. In early June 2019 the northern part was particularly wet, with larger areas covered with free water and quite high numbers of fen mire species like Savi’s Warbler *Locustella luscinioides*, Sedge Warbler, Bearded Tit *Panurus biarmicus*, Spotted Crake, Common Snipe and other waders. We thus conclude, that the complete absence of AW with geolocators in 2019 was not caused by unsuitable habitat conditions.

## 4. Results and discussion

### 4.1 General results: Performance of fieldwork and devices, recovery ratios, body condition of recovered birds

The catching of AW and fitting them with geolocators went very smoothly. In Servech, 29 adult males were captured within three days only. In Lithuania, 31 males were captured despite two days of weather unsuitable for mist-netting.

Recovery and recapture of AW in late May 2019 turned out to be more difficult than expected. This was probably due to a smaller team (smaller than previously planned) and a full day with bad weather in Servech. However, we probably found almost all of the returned geolocator birds, as we observed c. 25-30 singing males without geolocator and succeeded to recapture all 7 geolocator birds seen.

In Lithuania, one of us (Vytautas Eigirdas) already discovered and recaptured 10 geolocator birds in Alka polder previous to the arrival of the full team in late May. Thus, we were able to focus our activities on Tyrai marshes. Surprisingly, none of the observed 25-28 singing males carried a geolocator. The observation of a colour-ringed female and the detection and capture of a singing male ringed by us as juvenile bird in 2018 proofed our search to be performed thoroughly. The absence of geolocator birds in Tyrai marshes is amazing considering the high recovery ratio found in the nearby Alka polder.

In total, 32 % of GL birds were recovered, 24 % in Belarus and 39 % in Lithuania (Table 1). These values represent exactly the limits of the previously detected return ratios in adult AW of 25-40 % in Germany (Wawrzyniak & Sohns 1977) and Poland (Dyrcz & Zdunek 1993).

**Table 1:** Recovery ratios of AW fitted with geolocators.

| Study site | AW fitted with geolocators in 2018 | AW recaptured in 2019 and 2020 (=retrieved geolocators) | recoveries in % |
| --- | --- | --- | --- |
| Servech, Belarus | 29 | 7 | 24 % |
| *Alka polder, Lithuania* | 24 | 12 | 50 % |
| *Tyrai mire, Lithuania* | 7 | 0 | 0 % |
| total for Lithuania | 31 | 12 | 39 % |
| **total all sites** | **60** | **19** | **32 %** |

All the 19 retrieved geolocators contained full data sets for almost the entire migration cycle. The devices stored data from periods of 267 to 319 days; valid data logging stopped between 8^th^ April (after 267 days) and 30^th^ May (319 days), median was 12^th^ May (301 days).

There was no difference in size of the captured adult male AW in N-Belarus and Lithuania (Table 2). The slightly larger wing length and body mass of AW that were recovered after one year, compared to the total of captured AW males in 2018, was not statistically significant, but probably a first indication, that larger and stronger birds have a higher survival and return rate than the mean. The 19 geolocator birds recaptured in 2019 had significantly longer wings and higher body mass compared to the total of 61 birds captured in 2018 (Fig. 1 and 2). This is a clear indication that winter survival and return ratio is positively correlated with wing length and body mass of the birds.

**Fig. 1:**
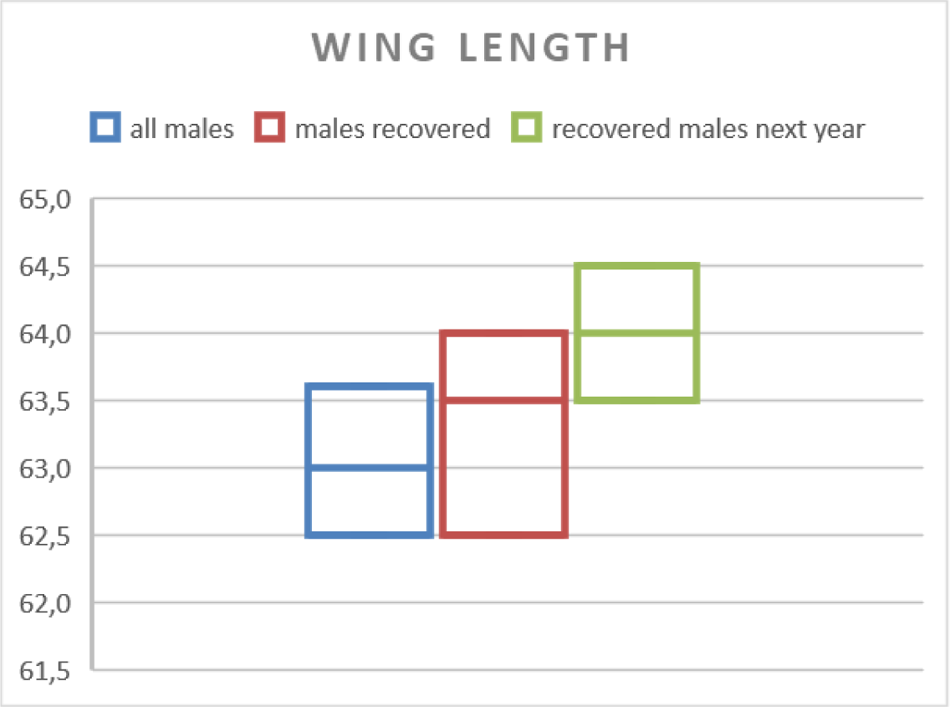
Median and 95% confidence interval of wing-length (in mm) of all adult males captured in 2018 (n = 61), those males recovered later, measurements from 2018 (n = 19) and the measurements of the recovered males in 2019 (n = 17).

**Fig. 2:**
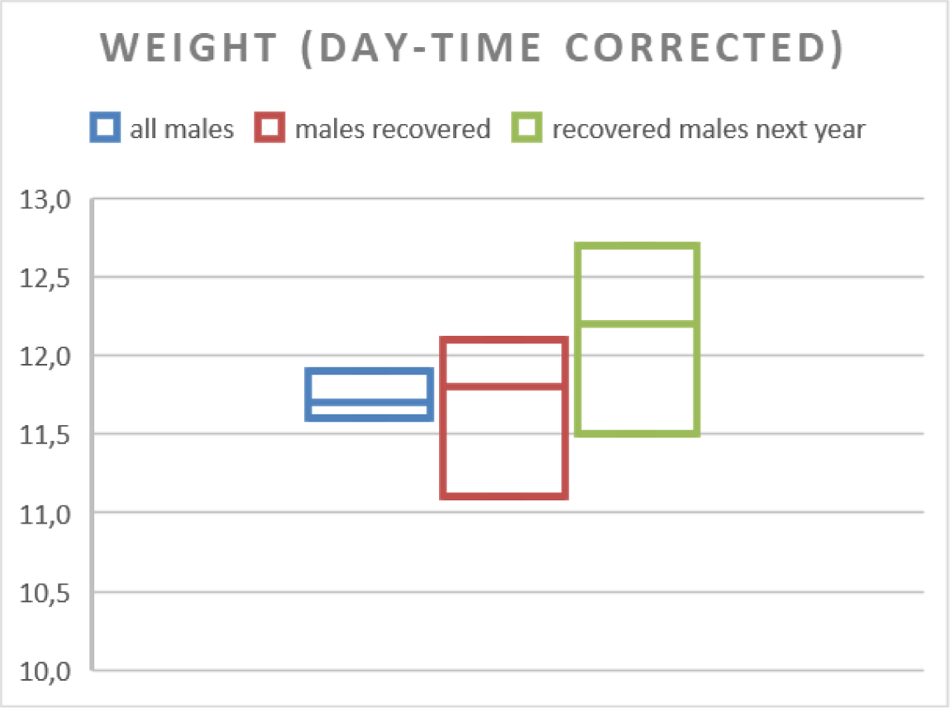
Median and 95% confidence interval of body mass (daytime-corrected weight in g) of all adult males captured in 2018 (n = 61), those males recovered later, measurements from 2018 (n = 19) and the measurements of the recovered males in 2019 (n = 17).

**Table 2:** Wing length (mm) and weight (g, day-time corrected) of adult AW males captured in Belarus (Servech) and Lithuania (Alka polder and Tyrai) in 2018.

|  | Belarus (n=29) |  | Lithuania (n= 32) |  |
| --- | --- | --- | --- | --- |
|  | wing | weight | wing | weight |
| upper CI | 64,0 | 12,0 | 63,5 | 11,9 |
| **median** | **63,5** | **11,7** | **63,0** | **11,8** |
| lower CI | 62,3 | 11,3 | 62,4 | 11,6 |
| **mean** | **63,3** | **11,7** | **63,2** | **11,8** |

### 4.2 Migration routes, wintering region and annual cycle of the 19 recovered Aquatic Warblers

There was no significant difference in migration routes and wintering sites of Lithuanian and NBelarusian birds. Fourteen or fifteen birds overwintered in Mali (6 from Servech, 8-9 from Alka polder), 9 of them in the region of the Inner Niger Delta (IND), the other 5-6 in SW-Mali or along the border to Burkina Faso (probably Sourou river floodplain). Only one bird did spend the winter in the Senegal delta (BP 343, from Servech), one in SE-Mauritania (BP 308, from Alka polder), one or two birds wintered in northern Burkina Faso (BP 303 and 319, both from Alka polder) and another bird in northern Nigeria (BP 300, from Alka polder).

At least 12 of the birds performed a distinct loop migration. In seven, the autumn and spring migration routes were completely different. Autumn migration was performed more or less along the Atlantic Sea coast via France, Spain, Portugal and Morocco, whereas spring migration was performed more directly with regular crossings of the Mediterranian Sea and/or stopovers along the Mediterranean coast of Algeria, Spain and France, on Mallorca island, on Sicily, in central and northern Italy.

On autumn migration, many birds had short stopovers of one or few days in Poland (5 birds), Germany (11), Czech Republik (3), The Netherlands (1), Luxembourg (1) and France (15), stopovers of at least 3 days were performed mainly in France, long stopovers with most likely fueling up in Spain (12), Portugal (8), Morocco/West Sahara (4) and Mauritania (5). Four birds had stopovers of several days in Senegal. Exceptions are two birds that wintered in Burkina Faso with stopover in Mali (BP 303), and in Nigeria (BP 300) with stopover in Algeria. The latter bird had the most extraordinary migration route: Coming from Alka polder in Lithuania, it had a very long stopover in the region of Gibraltar, then stopped in front of the Atlas mountains, passed them finally at the eastern side and flew non-stop to Nigeria and wintered there successfully. On its way back, it had a long stopver at the Mediterranean coast of NE-Algeria and returned then, probably within a few days (but no valid data), to Lithuania.

On spring migration, many birds had long stopovers in Mauritania (1 bird), Morocco (7) at the Algerian coast (3), two more at the southern coast of Spain (probably Cota Doñana). Short stopovers of one or few days were performed, besides Mauritania and Morocco, in Spain (6), France (5) and Belgium (1) on the western flyway, and Algeria (5), Mallorca (1), central (2) and northern (1) Italy, Croatia/Serbia (1) and Czech Republik (1) on the more easterly route. Several birds had very short stopovers in Germany (8) and three others also in NE-Poland and Russia/Kaliningrad region.

Summarising the data of the 19 recovered geolocator birds, adult AW males spent more than half of the year (184 days on average) at the wintering sites, c. 50 days each on autumn and spring migration, and only 81 days at the breeding sites (Fig. 3). On autumn migration, main stopovers were in Spain (16 days on average), and on spring migration in Morocco, N-Algeria and S-Spain (28 days on average) (Fig. 3, Table 4).

**Fig. 3:**
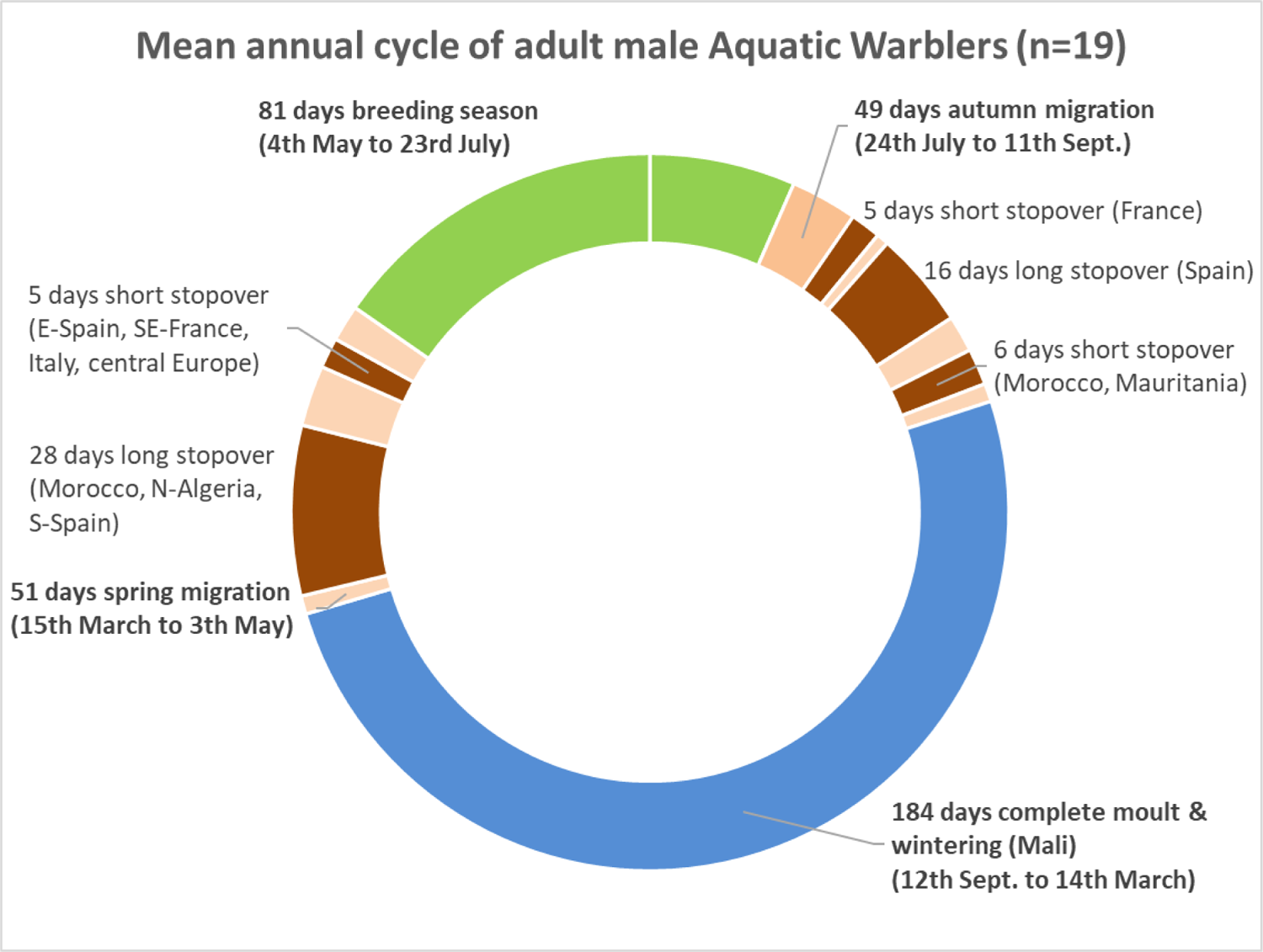
Mean annual cycle of adult male Aquatic Warblers according to the data of the 19 gelocator birds of this study. Mean values of dates and duration of stopovers and the countries hosting the majority (>70 %) of birds are indicated.

**Table 4:** Overview over the stopover (>2.5 days) and wintering sites country by country, in the order of migration.

| **Overview on AW stopover (>2.5 days) and wintering sites (19 adult males)** |  |  |  |  |  |  |  |  |  |
| --- | --- | --- | --- | --- | --- | --- | --- | --- | --- |
| migration period | Country | number of stopover sites | number of birds | duration of stay (days) |  | period of occurrence |  | median arrival | median departure |
|  |  |  |  | mean | median | from | until |  |  |
| autumn | Germany | 1 | 1 | 3,0 | 3 | 15.08. | 18.08. | 15.08. | 18.08. |
|  | Luxembourg | 1 | 1 | 4,0 | 4 | 14.08. | 17.08. | 14.08. | 17.08. |
|  | France | 6 | 10 | 6,9 | 6 | 22.07. | 31.08. | 06.08. | 12.08. |
|  | Spain | 10 | 12 | 15,8 | 13 | 26.07. | 13.09. | 10.08. | 25.08. |
|  | Portugal | 4 | 8 | 8,5 | 8 | 06.08. | 14.09. | 21.08. | 27.08. |
|  | Morocco | 4 | 6 | 10,6 | 8 | 10.08. | 22.09. | 09.09. | 17.09. |
|  | Mauritania | 5 | 5 | 10,3 | 9 | 19.08. | 23.09. | 11.09. | 19.09. |
|  | Senegal | 4 | 4 | 5,8 | 6 | 22.08. | 09.09. | 27.08. | 31.08. |
| wintering | Senegal | 1 | 1 | 202,0 | 202 | 25.08. | 15.03. | 25.08. | 15.03. |
|  | Mauritania | 1 | 1 | 198,0 | 198 | 26.09. | 11.04. | 26.09. | 11.04. |
|  | Mali | 15 | 14 | 196,4 | 187 | 27.08. | 12.04. | 11.09. | 11.03. |
|  | Burkina Faso | 2 | 2 | 173,5 | 174 | 04.09. | 03.03. | 07.09. | 27.02. |
|  | Nigeria | 1 | 1 | 157,0 | 157 | 26.09. | 02.03. | 26.09. | 02.03. |
| spring | Mauritania | 3 | 2 | 9,7 | 3 | 05.03. | 06.04. | 10.03. | 12.03. |
|  | Morocco | 11 | 10 | 18,7 | 16 | 17.02. | 12.04. | 11.03. | 07.04. |
|  | Algeria | 6 | 6 | 18,8 | 14 | 07.03. | 24.04. | 25.03. | 16.04. |
|  | Spain | 6 | 6 | 18,2 | 5 | 16.02. | 19.04. | 13.04. | 17.04. |
|  | Spain/France | 1 | 1 | 7,0 | 7 | 09.04. | 16.04. | 09.04. | 16.04. |
|  | Italy | 2 | 2 | 6,0 | 6 | 06.04. | 25.04. | 14.04. | 20.04. |
|  | Croatia/Serbia | 1 | 1 | 5,0 | 5 | 26.04. | 01.05. | 26.04. | 01.05. |
|  | Germany | 1 | 1 | 3,0 | 3 | 21.04. | 24.04. | 21.04. | 24.04. |
|  | Poland | 2 | 2 | 3,5 | 4 | 27.04. | 04.05. | 30.04. | 03.05. |
|  | Russia-Kalining | 2 | 1 | 9,5 | 10 | 29.04. | 19.05. | 03.05. | 12.05. |
|  | Belarus | 4 | 2 | 4,4 | 4 | 09.05. | 24.05. | 14.05. | 16.05. |

**Table 5:**
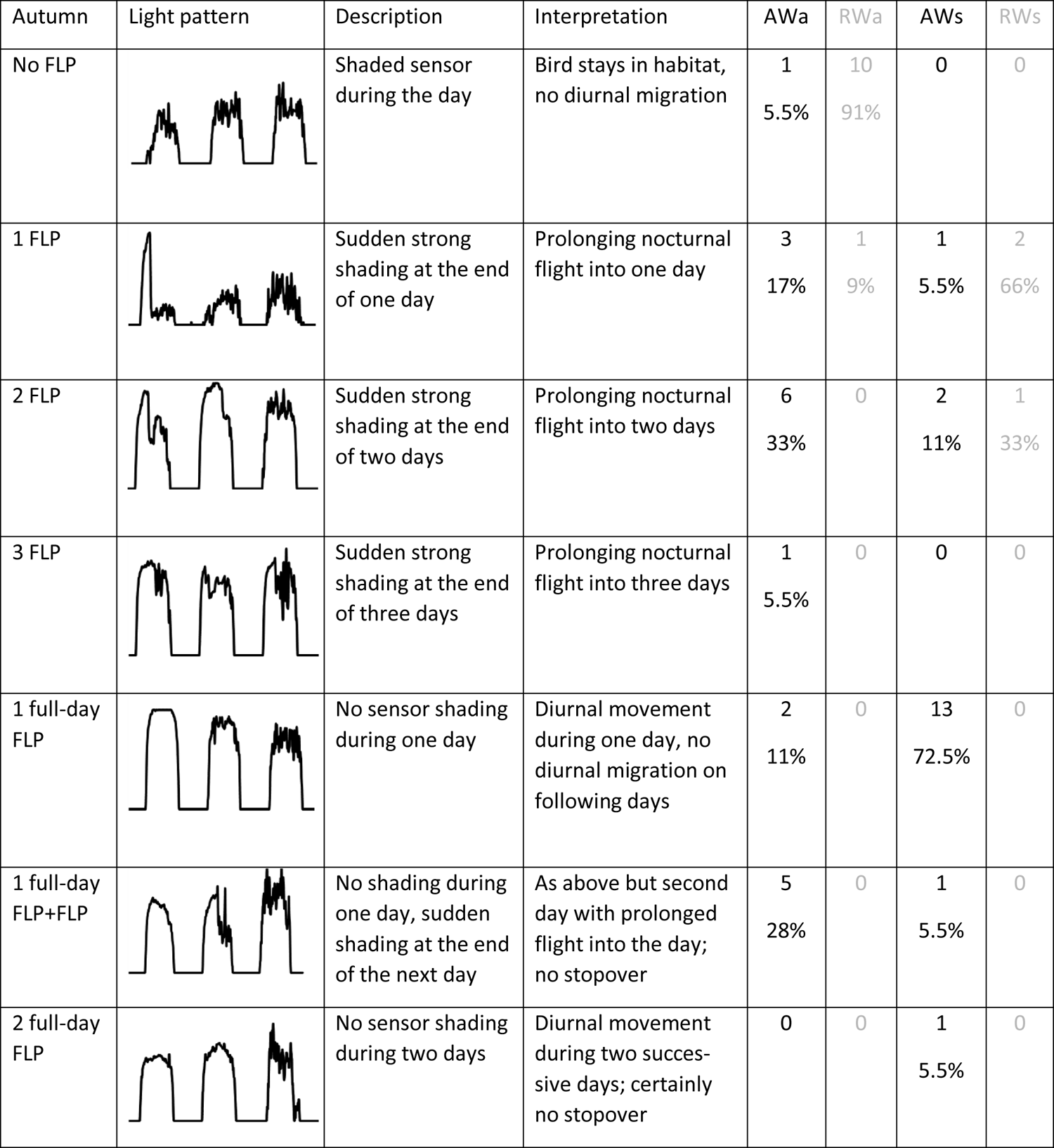
Detailed autumn and spring overview of daytime light pattern anomalies (FLP) recorded by geolocators while barrier crossing. Each category is accompanied by a representative figure of recorded light intensities, description of the anomaly and the most plausible interpretation, numbers of individuals. Specific % of occurrence, in line with previous publications^8^. AW = Aquatic Warbler, RW = Reed Warbler (from Jiguet et al. 2019); a for autumn, s for spring.

| Autumn | Light pattern | Description | Interpretation | AWa | RWa | AWs | RWs |
| --- | --- | --- | --- | --- | --- | --- | --- |
| No FLP |  | Shaded sensor during the day | Bird stays in habitat, no diurnal migration | 1<br>5.5% | 10<br>91% | 0 | 0 |
| 1 FLP |  | Sudden strong shading at the end of one day | Prolonging nocturnal flight into one day | 3<br>17% | 1<br>9% | 1<br>5.5% | 2<br>66% |
| 2 FLP |  | Sudden strong shading at the end of two days | Prolonging nocturnal flight into two days | 6<br>33% | 0 | 2<br>11% | 1<br>33% |
| 3 FLP |  | Sudden strong shading at the end of three days | Prolonging nocturnal flight into three days | 1<br>5.5% | 0 | 0 | 0 |
| 1 full-day FLP |  | No sensor shading during one day | Diurnal movement during one day, no diurnal migration on following days | 2<br>11% | 0 | 13<br>72.5% | 0 |
| 1 full-day FLP+FLP |  | No shading during one day, sudden shading at the end of the next day | As above but second day with prolonged flight into the day; no stopover | 5<br>28% | 0 | 1<br>5.5% | 0 |
| 2 full-day FLP |  | No sensor shading during two days | Diurnal movement during two successive days; certainly no stopover | 0 | 0 | 1<br>5.5% | 0 |

### 4.3 Main stopover sites in autumn (Annex 1, Table 4, Fig. 4)

Due to long-lasting bird ringing programmes in Belgium, Luxembourg, France, Spain and Portugal, many AW stopover sites are known in SW Europe (summary see Le Nevé et al. 2018). The 19 male AW with geolocators left their breeding sites on average on 24^th^ July and migrated more or less non-stop to France. This first migration phase was only interrupted by short stopovers of less than 2.5 days. Only one bird (BP 326) had a 3 days stopover in NW-Germany in the state of Schleswig-Holstein near the west coast, probably at the ‘Beltringharder Koog’, a coastal freshwater and salt marsh.

**Fig. 4:**
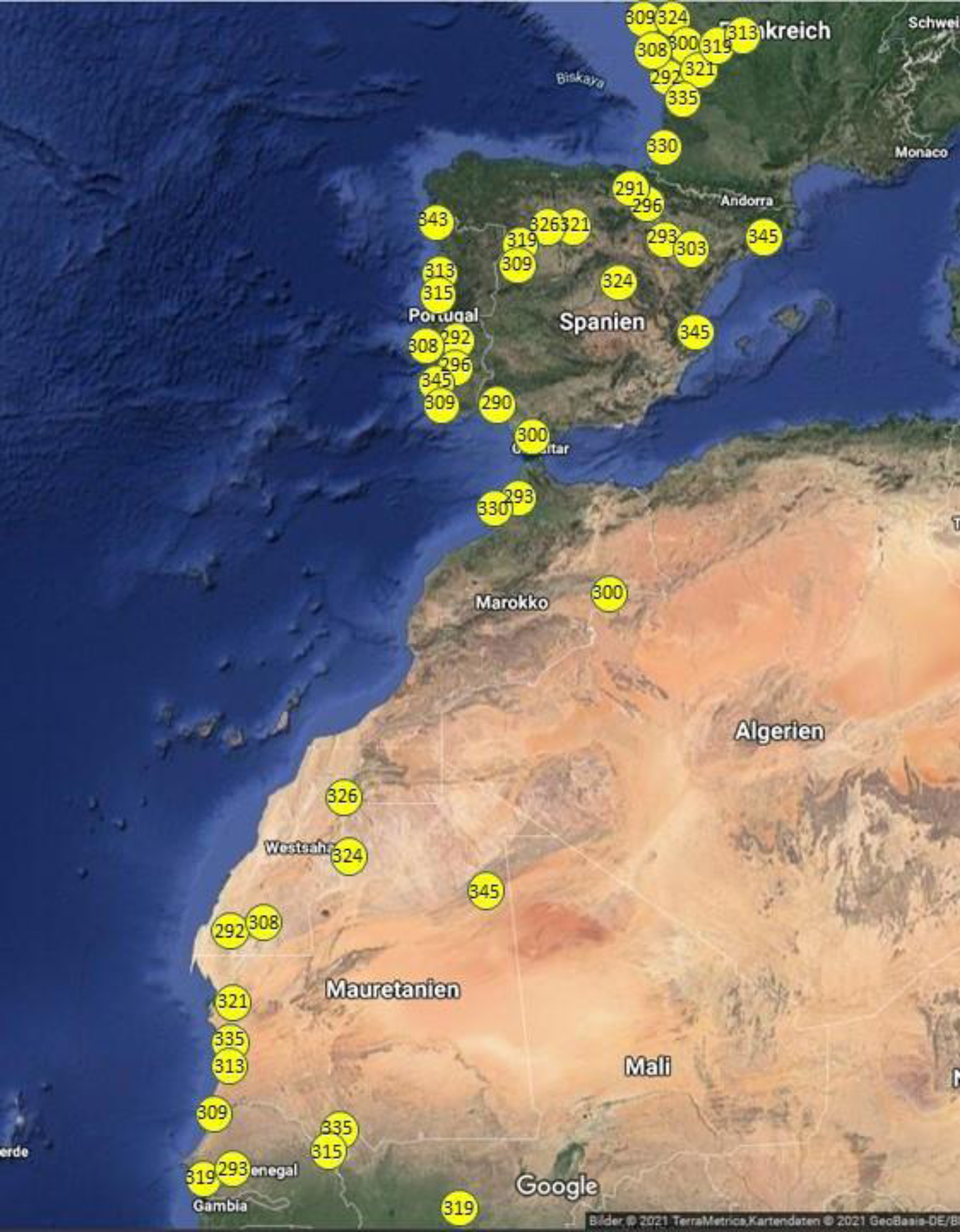
Approximate stopover locations (>2.5 days) on autumn migration of the 19 Aquatic Warblers; the ID number of the geolocator is indicated.

Ten birds had their first longer stopover of 4-16 days (mean 6.9 days) in the period from 22^nd^ July to 20^th^ August at the French Atlantic Sea coast (Gulf of Biskaya) from the Loire estuary southwards (Fig. 4, Annex 3). Two of the birds might have rested at the Loire estuary, five more at the Garonne estuary.

The main stopover sites on autum migration with a durance of stay of mostly 11-29 days, mainly between 10^th^ and 27^th^ August, are located on the Iberian peninsula in the northern half of Spain and along the Portuguese and South-Spanish Atlantic Sea coast (Fig. 4). In northern and central Spain, two birds (BP 321, BP 326) possibly rested at the well-known Laguna de La Nava, three more (BP 293, 303, 345) somewhere along the Ebro floodplain and estuary. One bird could not be assigned to any known stopover site. Only one bird has rested at the Spanish Mediterranean coast (BP 345, Ebro delta and S of Valencia), and one bird each probably at the Cota Doñana (BP 290) and near San Fernando (Cadiz)(BP 300). All stopover sites in Portugal were presumably located at river estuaries and lagoons along the Atlantic Sea coast.

One bird (BP 293) had its first longer stopover of 20 days in NW-Morocco, probably at the well-known coastal lagoon of Merja Zerga. This bird had previously stayed in NE-Spain for 5 days only. Three birds had shorter stopovers of 5-8 days in Morocco and West Sahara, one bird (BP 292) stayed in West Sahara for 14 days, its longest stopover on autum migration, after having rested around the Garonne estuary for 5 days and at the Portuguese coast (probably Estuario do Sado) for 9 days.

Five birds had shorter stopovers of 4-9 days in Mauritania, three of them probably at wetlands at or near the coast (BP 313, 321, 335) and one (BP 335) within or near the middle Senegal floodplain. Only one bird (BP 309) rested for 6 days in the region of Djoudj/Senegal delta, two more birds (BP 293, 319) in the region of the Saloum river floodplain and delta, a fourth bird (BP 315) probably also in the middle Senegal floodplain.

One bird (BP 300 from Alka polder) performed a very special migration: After stopovers for 4 days in W-France and 25 days in S-Spain near Cadiz, it rested again for 5 days in the desert somewhere in the border region of NE-Morocco and Algeria (possibly at the large desert lake Djorf Torba in Algeria), and then crossed the central Sahara on a more easterly flyway directly to its wintering site in N-Nigeria. The full Sahara crossing (2,460 km) took 8 days with only one short interruption of 1.5 days (on way back this bird needed 5 days for 2,650 km non-stop crossing of the central Sahara to the Algerian coast).

Unfortunately, it is nearly impossible to assign the positions of resting geolocator birds to specific sites in the deserts of West Sahara and Mauritania by using the google maps satellite images, because they are mainly or exclusively taken in the dry season. To identify possible stopover sites with higher precision, high resolution images from exactly the time window when the geolocator birds were resting there are required, that means from mid-August to mid-September of the year 2018. This is because suitable stopover habitats in the desert may occur for short periods of a few weeks and disappear again according to the distribution of precipitation in exactly that year.

Summarising, autumn migration of the 19 studied adult AW males lasted 49 days on average, with a mean departure on 24^th^ July and mean arrival at the wintering sites on 12^th^ September. Nine birds had a first short stopover in France, and most birds had a longer stopover on the Iberian Peninsula (14 birds) and NW-Morocco (1 bird), one other bird not before reaching West Sahara. Fourteen birds had another shorter stopover of mostly 4-7 days S of the Sahara, mainly in Mauritania and Senegal, before reaching their wintering grounds in Mali and NW-Burkina Faso. One bird (BP 313) had no longer stopover at all, but rested only three times for 5-6 days at coastal sites of France, Portugal and Mauritania.

### 4.4 Wintering sites (Annex 2, Fig. 5)

In the southern Sahara and adjacent Sahel region, larger wetlands that are probably suitable for AW are situated almost isolated in the desert and are more or less well visible on satellite images. Many of them are known from targeted terrestrial search for AW in the period 2007-2012 or older accidental ring recoveries (summary see Tegetmeyer et al. 2018).

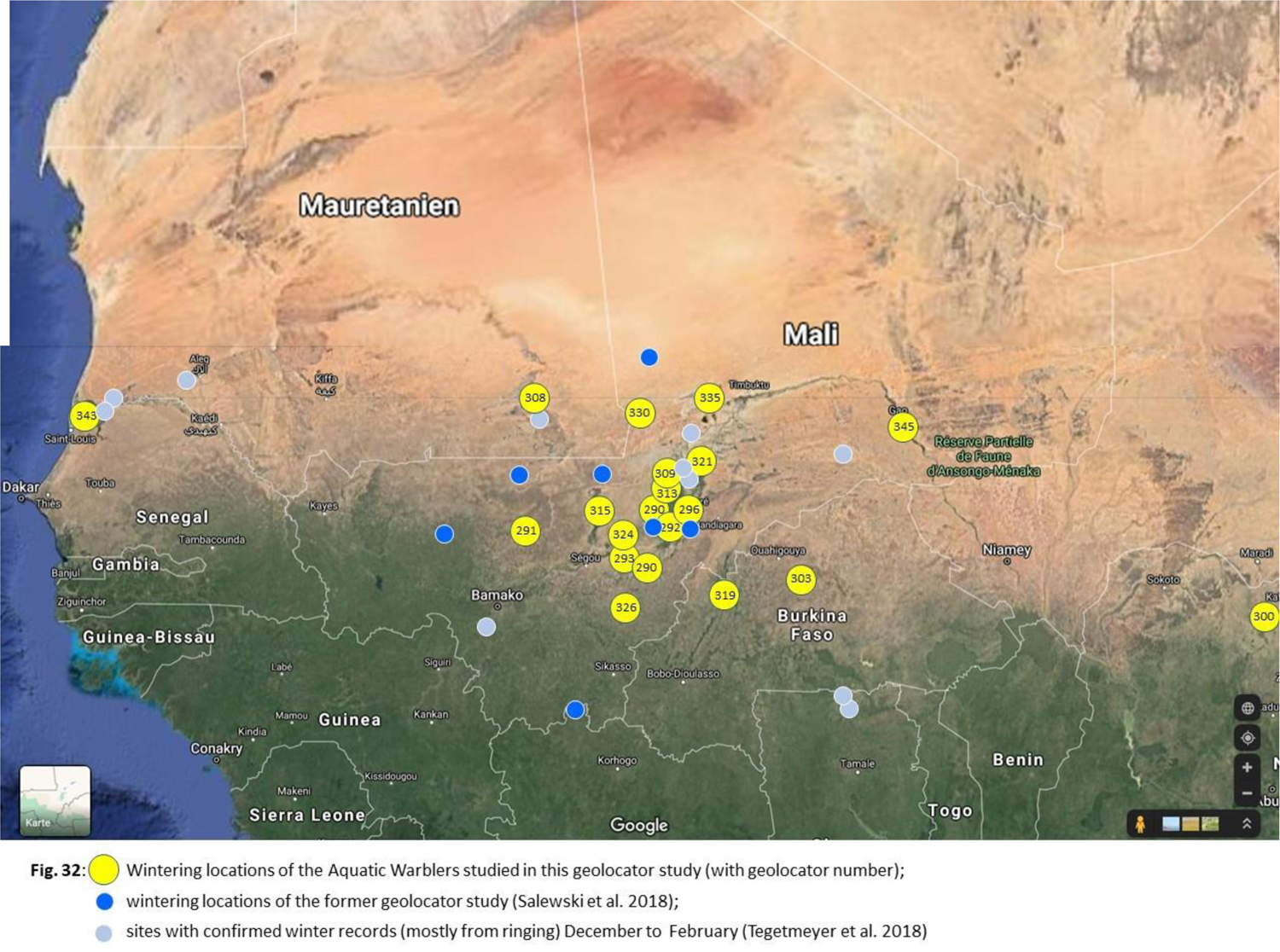

In contrast to our expectations, only one of the 19 AW overwintered in the Senegal delta (probably Djoudj; BP 343, from Servech). The majority of birds spent the winter in Mali (14-15) and adjacent parts of Mauritania (1) and NW-Burkina Faso (1-2, one of them in border region to Mali), one bird overwintered in N-Nigeria (Fig. 32). In Mali, 10 birds overwintered in the Inner Niger Delta (IND) (6) or nearby parts of the floodplains of Niger and Bani rivers. One more wintering site (BP 345) can be assigned to the Niger floodplain near Gao in E-Mali, one (BP 315) to the River Sahel floodplain W of the IND, and another one (BP 291) possibly to the Baoulé river floodplain (possibly Boucle du Baoulé National Parc) NNE of Bamako. The bird wintering in the SE-corner of Mauritania (BP 308) was probably located in the inland delta of the Issil river, not far away from the wintering site Adel Bagrou recorded in 1999 (Tegetmeyer et al. 2018). In Burkina Faso, one bird (BP 319) in the border region of Burkina and Mali might have wintered in the wide floodplain of Sourou river (but possibly in the Reserve de Bay on the Mali side of the floodplain), the other bird (BP 303) probably at one of the large fresh water reservoirs (dammed rivers) of northern central Burkina Faso.

In the IND, most or even all wintering AW were located in the outer reaches of the delta, but none in the central, higher flooded part. Only one bird (BP 290) changed the site during the wintering period. After 113 days at the south-western margin of the IND it shifted 90 km in SSW direction to the region of the Bani floodplain, where it spent 64 more days. All other birds (!) did spend the full winter (139-222 days, average 184 days) at just one location.

Altogether, the results of the geolocator study match quite good with former knowledge on the wintering sites from the former geolocator study on central Ukrainian and W-Belarusian birds (Salewski et al. 2019) and ringing activities in Senegal/Djoudj, the IND and S-Mauritania (Le Nevé et al. 2018, Flade et al. 2011, Poluda et al. 2012, Foucher et al. 2013). Most localisations of geolocator birds (15 out of 19) can be assigned with medium to high probability to known, confirmed wintering sites or larger areas with probably similar habitats.

### 4.5 Main stopover sites in spring (Annex 3, Fig. 6)

Stopover sites of AW in spring are far less well-known from bird ringing than those in autumn, because there exist much fewer ringing schemes in spring in general, and along the Mediterranean coast and in NW-Africa (Morocco, Algeria). For this reason, and the equinox problem as well, assignment of geolocator positions of resting AW to known specific sites is more vague in spring. This counts the more in comparison to the wintering sites, were the birds spend up to more than 200 days at one location just between the two equinox phases.

**Fig. 6:**
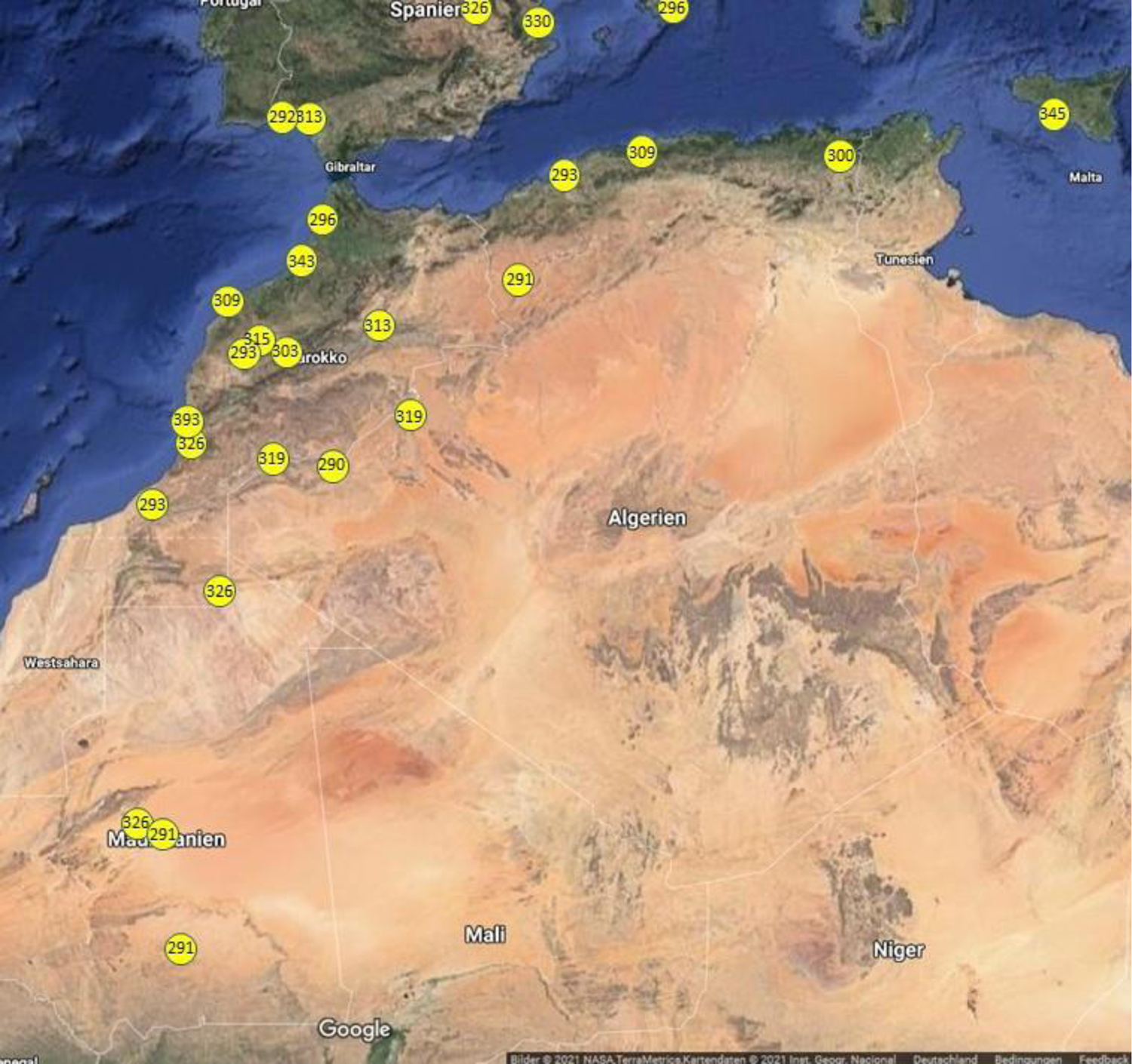
Approximate stopover locations (>2.5 days) in NW-Africa on spring migration of the 19 Aquatic Warblers; the number of the geolocator is indicated.

**Fig. 7:**
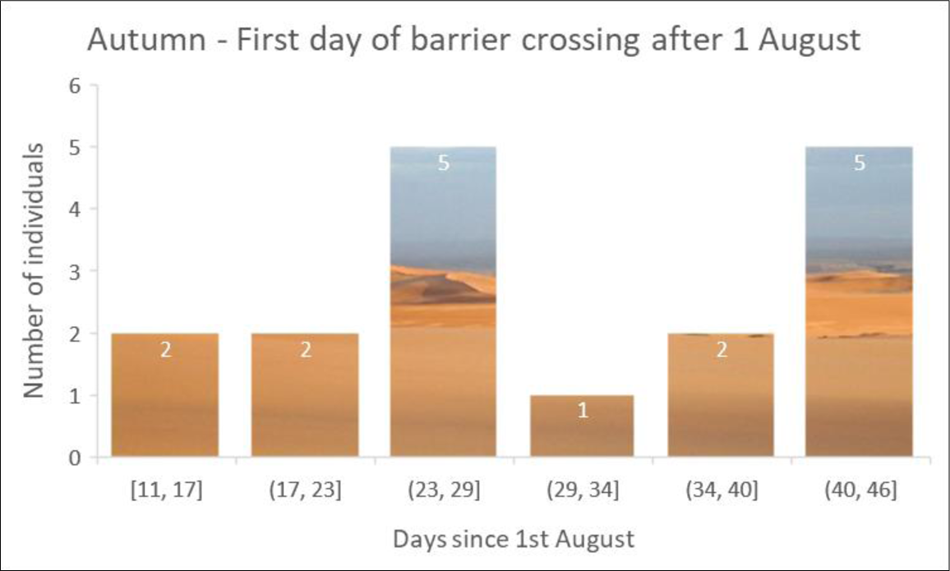
Phenology of the first day of detected full-light pattern in the 18 aquatic warblers in autumn, revealing two different ‘waves’. The reasons for these waves are unknown so far.

**Fig. 8:**
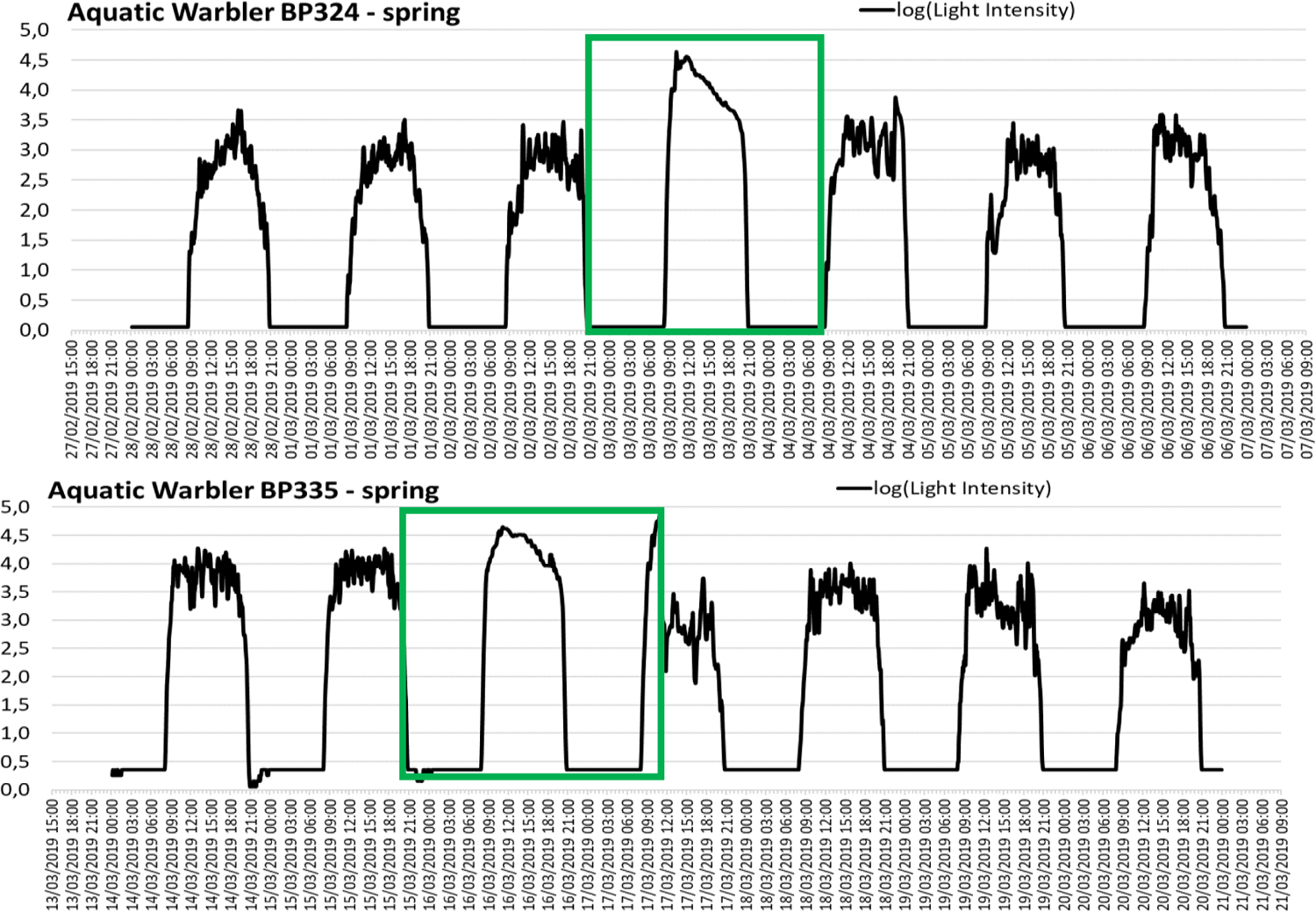
Light intensity (log-transformed) recorded by geolocators during spring desert crossing for two individuals. Green rectangles represent possible continuous flight bouts.

**Fig. 9:**
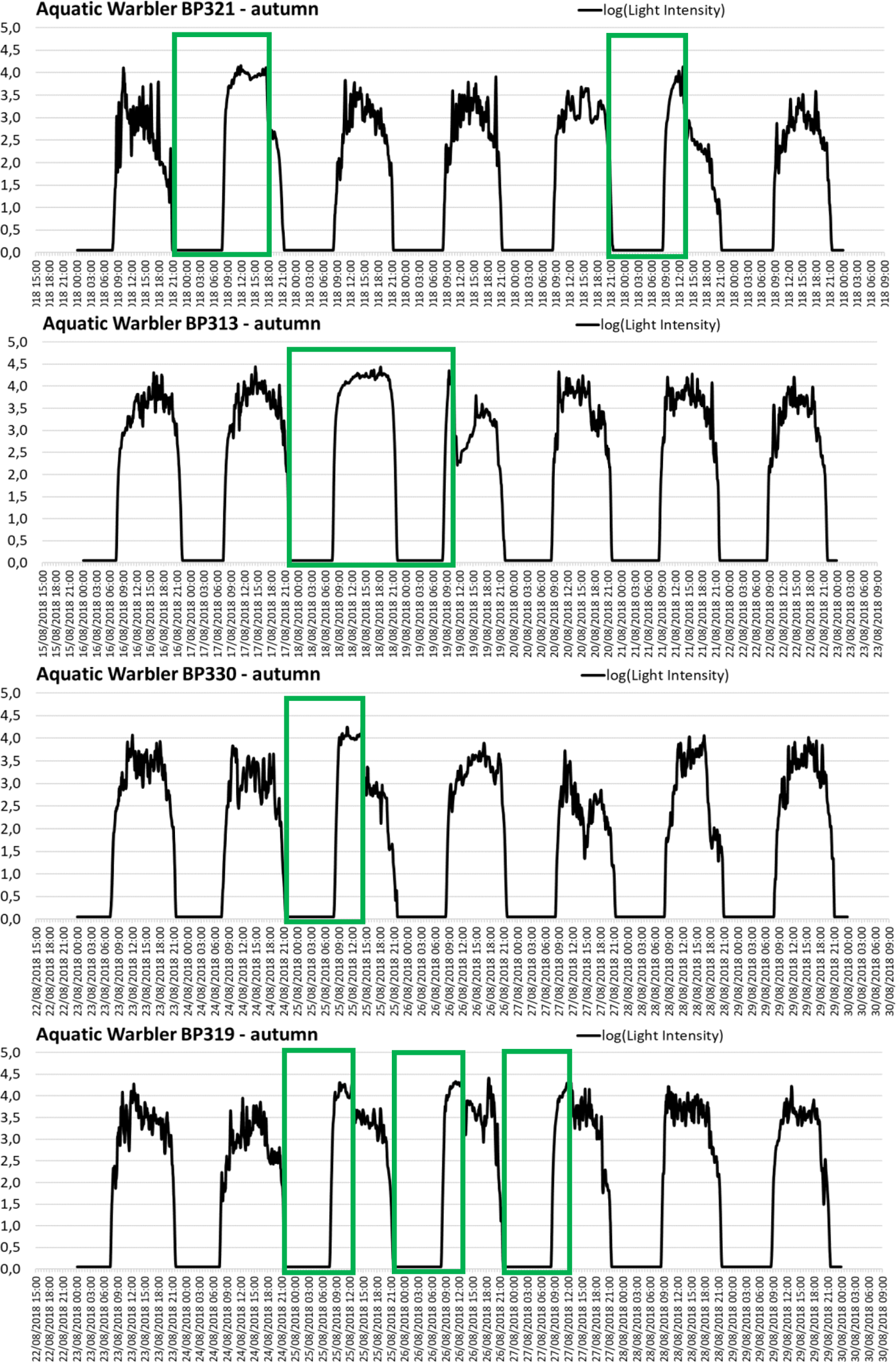
Light intensity (log-transformed) recorded by geolocators during autumn desert crossing for four individuals. Green rectangles represent possible continuous flight bouts.

**Fig. 10:**
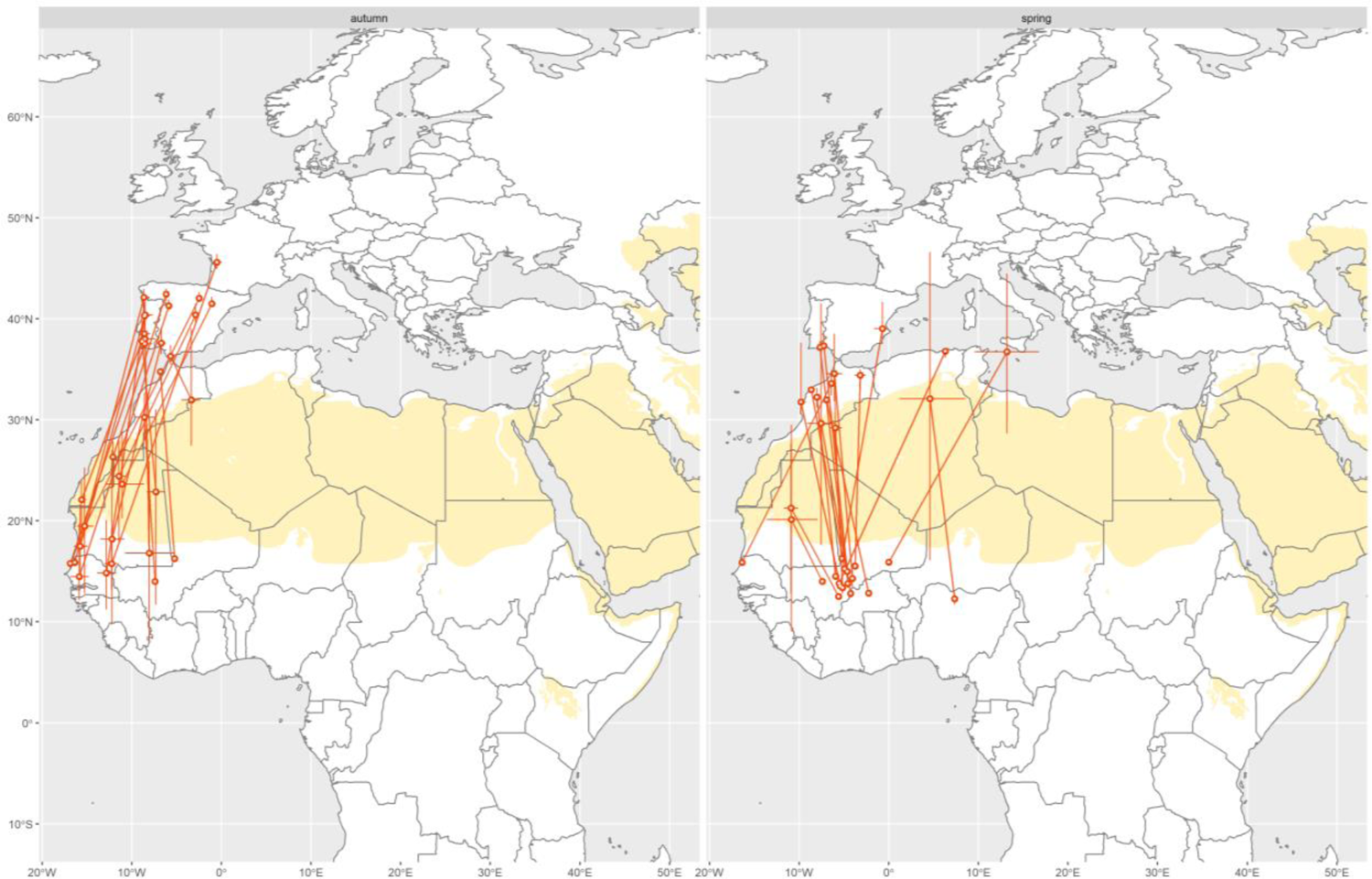
Stationary sites of individual birds (median ±2.5/97.5 percentiles of location estimates) just prior and after the occurrence of the full light pattern(s) detected during autumn and spring.

Spring migration started between 3^rd^ March (Nigeria) and 13^th^ April (Mali) and lasted 51 days on average. Mean departure date from the wintering grounds was the 15^th^ March. Most AW started the spring migration with instant Sahara crossing to reach their long spring stopover sites in Morocco and the coast of N-Algeria and S-Spain. Only six birds had short stopovers of 3-7 days in Mauritania and SMorocco/West Sahara before arriving at their main spring stopover. One bird (BP 291) left its wintering site in W-Mali on 7^th^ March to have a short (3 days) and then a long stopover (23 days) in the southern Sahara in Mauritania, jumped then over the Sahara to NW-Algeria (6 days stopover) before returning directly to the breeding site in Alka polder/Lithuania.

Morocco, the Mediterranean coast of Algeria und probably Cota Doñana in S-Spain are of major importance as resting areas on spring migration just after Sahara crossing and before the final jump to the central European breeding sites. Seven birds had long stopovers of 15-45 days in Morocco. BP 326 split its main spring stopover in 18 days in West Sahara (possibly at a small desert lake) and 13 days at the Atlantic Sea coast S of Agadir. BP 293 divided its spring stopover even in three periods: 32 days in SE-Morocco (Wadi Draa region), 7 days probably in the Sous Massa NP S Agadir and 11 more days somewhere N of Marrakesh. After another short stopover of 5 days in N-Algeria it jumped directly back to its breeding ground in Lithuania. BP 309 divided its long spring stopover into 15 days SW El Jadida in Morocco and 18 days in N-Algeria.

The AW wintering in Nigeria (BP 300) started its spring migration instantly with a non-stop flight of 2,650 km over the central Sahara (interrupted only by short rests during daytime) to the Algerian coast, probably in the region of Lac Fetzara. It needed 5 days with three stopovers during day and one full day migration to manage this travel (530 km per day on average).

One important reason for very long spring stopovers in Morocco/Algeria could be, that the AW are forced to leave their wintering sites earlier than necessary for migration because the wintering habitats are drying up and thus becoming increasingly unsuitable for the birds. If this would be the case, these long stopovers could also be regarded as the last stage of wintering.

Most important stopover and refuelling sites in Morocco are, according to our geolocator data, probably mainly situated along the Atlantic coast and match roughly with known sites of AW occurrence (Merja Zerga SSW Larache) or with confirmed suitable habitats (Souss Massa NP S Agadir, Ramsar site Sidi Moussa–Oualidia SW El Jadida, lower Sibou river and Lac Sidi Boughaba NP). Some – according to geolocator data – inland resting sites in inland Morocco (N and NE Marrakesh and along the border to Mauritania/Wadi Draa region) and Algeria can hardly be assigned to specific sites without analysing satellite images from exactly the time window of AW migration (early March to mid-April 2019). Additionally, geolocator localisations are especially imprecise around equinox (21^st^ March), making the assignment to specific sites even more difficult.

In Algeria, long stopovers of 18-48 days have been detected in the West near the border to Morocco (1 site) and near the Mediterranean coast (2 sites). Two AW had long stopovers of 41 und 53 days, respectively, in S-Spain, probably in the Cota Doñana National Parc.

After these long refuelling stopovers in Morocco/Algeria and S-Spain, AW travelled quickly back to their breeding grounds, with stopovers of not more than 3-8 days (average 5 days). These short stopovers were located in E-Spain (BP 326, 330), on Mallorca (BP 296), on Sicily (BP 345), at the Spanish/French Mediterranean coast (BP 319), in central Italy (BP 324), in the region of Save and Danube floodplains in Croatia/Serbia/Bosnia (BP 324) and in western Germany (North Rhine-Westphalia, BP 330).

Reaching central Europe, most AW returned directly to their breeding grounds. Only five birds visited other sites in central Europe for a few days before arriving at their tagging and recovery place: Two birds stopped for three days in the Mazury Lake district (BP 330, going to Alka polder) and in the middle Biebrza basin (BP 326, going to Servech). BP 290 had four stopovers in Vygonoshchanskoye (southern central Belarus), S of Minsk and in NW-Belarus before returning lately, on 25^th^ May, to Servech. BP 324 (going to Servech) stopped for 7 days probably at the Bjerezina floodplain (Bjerezinski Biosphere Reserve), and BP 315 visited probably the Pregolia floodplain and the Vistula Lagoon SW Kaliningrad before returning to Alka polder.

However, six geolocators stopped data logging already in Mali (BP 335), Algeria (BP 293, 300 and 309), Morocco (BP 313) and Italy (BP 345) because batteries were exhausted, so that the last phase of the travel of these birds could not be tracked.

### 4.6 Barrier crossing strategies

We detected a full daytime light pattern (FLP) in 17 of 18 autumn (94%) and in all 18 spring (100%) recorded barrier crossings for the AW, which occurred in one or two days (data of a 19^th^ bird BP 308 arrived too late for this analysis). This allows the identification of potential diurnal migration.

In autumn, a diversity of strategies consisted mostly in short FLP in one or two mornings (n=9, 50%), or a full-day FLP followed (n=5, 28%) or not (n=2, 11%) by a short morning FLP. For all incomplete FLPs, the average time spent in flight during daylight was 350 minutes after sunrise (± 137 s.d., median 345 min, range 80-600 min; n=24).

In spring, FLP was detected for one day for 14 individuals (78% of all individuals) and two days for 4 individuals (22%). For incomplete FLPs, the time spent in flight during the daylight was 363 minutes after sunrise (± 163 s.d., median 375 min, range 170-570, n=6). FLP during a complete day was detected for 15 individuals (including one individual with two full FLPs), representing 83% of individuals and 73% of the spring detected FLP. The main spring strategy of barrier crossing appears as a continuous flight during at least two nights and the day in between.

We produced a map reporting the estimates of stopover locations before and after the detected FLP(s). In autumn, two thirds concern tracks starting in Iberia and France, so that barrier crossing could concern either sea or desert crossing. However, almost all tracks include a complete or a part of the Sahara Desert, and for some individuals the FLPs occurred just before a stopover south of the Sahara, so we consider that the detected FLPs occurred during desert crossing. In spring, only 4 of 17 plotted tracks include the crossing of the Mediterranean or the Strait of Gibraltar, so again FLPs highly probably occurred during desert crossing, not sea crossing.

The strategy of the Aquatic Warbler to cross desert differs from that of Common Reed Warblers *A. scirpaceus* studied previously, which very rarely perform FLP in autumn, and only short FLP in spring – but no complete full-day FLP. Here we found mainly complete full-day FLP in spring, contrary to Adamik et al. (2016). Bibby & Green (1981) have also convincingly shown that Reed Warblers tend to migrate in shorter steps during autumn migration than the Sedge Warbler (which seems to be more similar to the AW in this aspect).

## 5. Summarising discussion

The study revealed expanded insights in the annual cycle of the AW and delivered a detailed picture of migration routes, migration and refuelling strategy, stopover system, desert crossing and wintering sites of the north-central European AW populations. The state of knowledge on the species summarised by Tanneberger & Kubacka (2018) has been significantly enlarged. The results especially on the importance of suitable stopover sites in Morocco/Algeria during spring migration just after Sahara crossing are groundbreaking with respect to the development of future conservation strategies and priorities for action.

Basis for these results was accurately executed and fine-tuned fieldwork on one hand and excellent geolocator devices without any malfunction on the other. When recapturing the birds, all birds were in a very good condition (see Table 1, Fig. 1 and 2). The total recovery rate of 32 % can be treated as excellent. However we cannot explain the complete absence of geolocator birds in Tyrai in 2019, although a high number of singing AW was present, and despite in the neighbouring site Alka polder we recovered 50 % of the birds equipped with geolocators in the previous year.

All 19 retrieved geolocators contained valid data for 267-319 days each, on average 301 days (c. 10 months, as announced in the Migrate Technologies instructions). This was sufficient to track autumn migration and to identify the wintering grounds of all recovered birds, and to track the spring migration for most of the birds. The assignment of the geolocator localisations to known sites of occurrence or larger areas of suitable habitat rendered to be difficult, especially for ephemeral habitats in the Sahara Desert and around equinox, and particularly in spring. Nevertheless, for 94 out of 115 stopover sites (>2.5 days rest) we could at least derive an “informed guess” of identification of possible or probable sites (overview see Table 8). Considering the previously known stopover sites and their distance to our geolocator localisations we estimate the accuracy of geolocator localisations at mostly better than +/80 km, sometimes probably only +/-150 km.

**Table 6:**
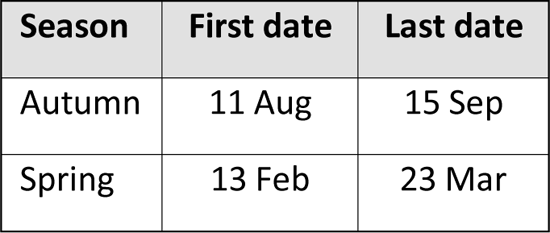
First and last dates with detected FLP (e.g. prolonged nocturnal flight into the day) while barrier crossing, by season. Very early barrier crossing compared to other European passerines, in autumn as in spring.

**Table 7:**
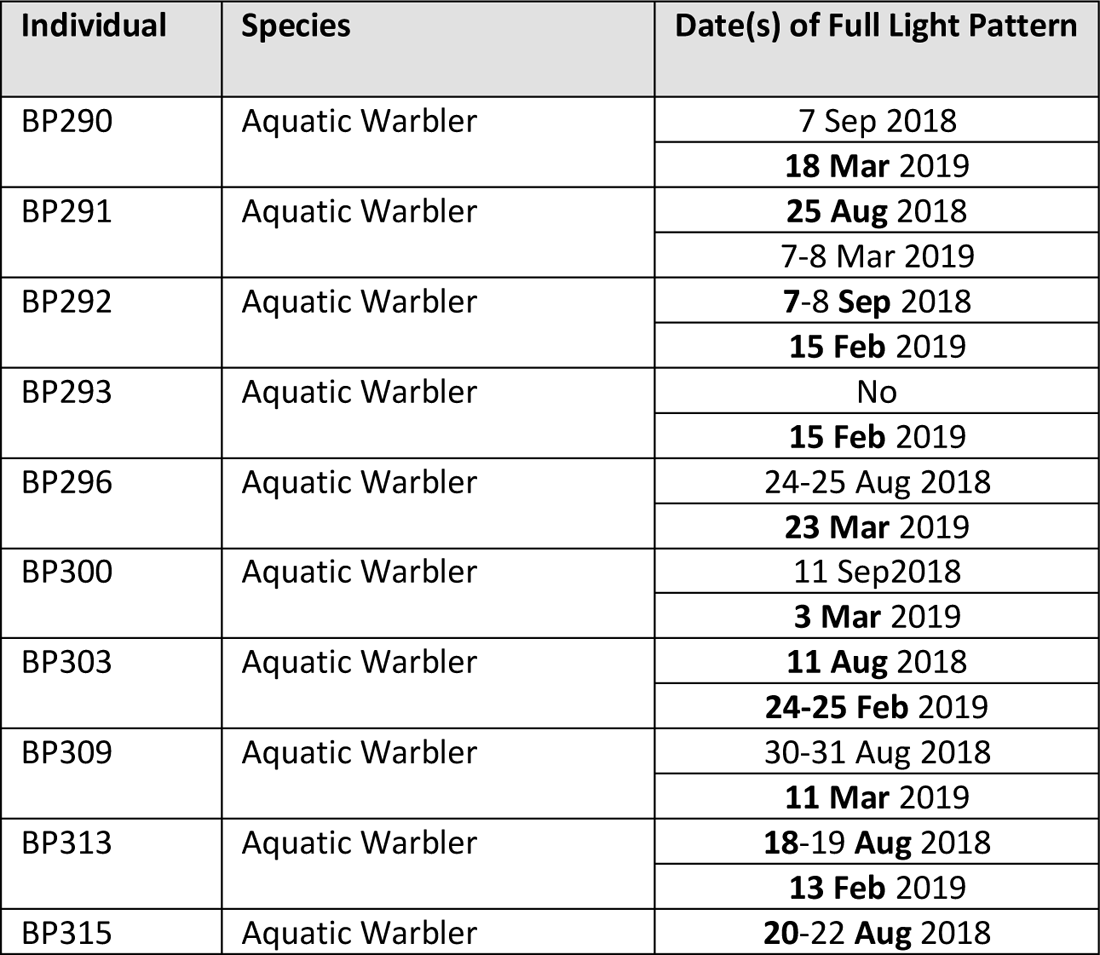
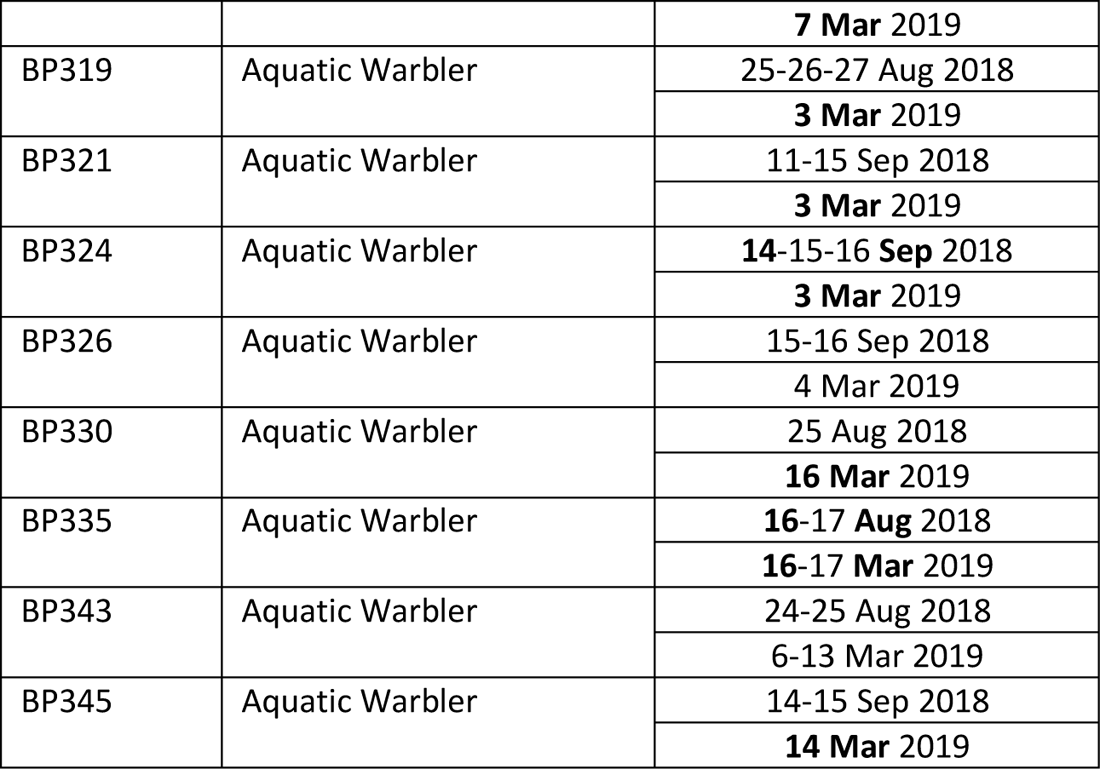
Tracked individuals, with references to logger number, species, breeding country, desert crossed during migration and dates with FLP(s) (in bold if a full-day FLP).

**Table 8:** Probability of correct assignment of geolocator localisations to known Aquatic Warbler stopover sites (e.g. ringing sites) or potentially suitable sites (=summary of annexes 3-5)

|  | Probability of correct appointment |  |  |  |
| --- | --- | --- | --- | --- |
|  | high | medium | low | no |
| **autumn migration** | **11** | **21** | **13** | **6** |
| Germany |  | 1 |  |  |
| Luxembourg |  | 1 |  |  |
| France |  | 6 | 3 | 1 |
| Spain | 4 | 5 | 4 |  |
| Portugal | 5 | 2 |  | 1 |
| Morocco | 2 | 1 | 1 | 2 |
| Mauritania |  | 2 | 2 | 2 |
| Senegal |  | 3 | 1 |  |
| Mali |  |  | 1 |  |
| Burkina Faso |  |  | 1 |  |
| **wintering** | **9** | **9** | **3** | **2** |
| Senegal |  | 1 |  |  |
| Mauritania |  | 1 |  |  |
| Mali | 6 | 6 | 2 | 1 |
| Burkina Faso |  | 1 | 1 |  |
| Nigeria |  |  |  | 1 |
| **spring migration** | **5** | **12** | **11** | **13** |
| Mauritania |  |  | 2 | 1 |
| Morocco | 1 | 6 | 2 | 3 |
| Algeria |  | 1 | 4 | 1 |
| Spain | 1 | 3 | 1 | 1 |
| Italy |  |  |  | 2 |
| France |  | 1 |  |  |
| Croatia | 1 |  |  |  |
| Germany |  |  |  | 1 |
| Poland | 1 |  |  | 1 |
| Belarus | 1 |  | 1 | 3 |
| Russia-Kaliningrad |  | 1 | 1 |  |
| **Whole annual cycle** | **25** | **42** | **27** | **21** |

One of the most important results is the fact, that 17 out of 19 AW overwintered in Mali and adjacent Burkina Faso and Mauritania, and only one in Senegal/Djoudj, where more were expected to hibernate, and one more in Nigeria. The huge importance of Mali with the Inner Niger Delta and surrounding floodplains of Baoulé, Sahel, Bani, and Sourou rivers for the survival of the species became obvious. However, the Senegal delta (Djoudj) could still have high importance for the – not yet studied Polish AW population.

As previously assessed (summarised by Le Nevé et al. 2018), the Iberian Peninsula has outstanding importance as stopover and refuelling area of adult male Aquatic Warblers in autumn. In spring, Morocco, S-Spain and the Mediterranean coast of Algeria, with average resting durance of 28 days, seems to be most important for these essential ecological requirements. This justifies to increase efforts to identify these sites in Morocco and Algeria on the ground and secure adequate protection.

It was possible to reveal the Sahara crossing strategy of the birds in detail by sophisticated data analysis of the full daylight pattern. This strategy is quite different to that of Reed Warbler and more similar to Sedge Warbler (Adamik et al. 2016, Bibby & Green 1981). Due to regular daily non-stop flights and Sahara crossings within a few days the AW relies much more on suitable stopover sites at the northern margin of the Sahara than other *Acrocephalus* species.

Last not least, it was now possible to derive the complete annual cycle of adult male AW from Lithuania and N-Belarus (Fig. 1). Especially impressive is the finding, that the AW spend more than half of the year, 184 days on average, in the wintering grounds at almost the same site (only one bird switched to another wintering site one time). Another c. 100 days they are on migration, and only c. 80 days at the breeding grounds.

However, all these results are only valid for adult AW males. The temporal and spatial proportions are likely to be quite different in females and juveniles (Wojczulanis-Jakubas et al. 2017, Le Nevé et al. 2018). Adult females leave the breeding grounds later on average, as c. 50 % do perform a second brood, which fledges not before late July, mostly probably until mid-August (Dyrcz et al. 2018). Because females care about their young without the support of males, we assume, that females leave the breeding grounds 2-3 weeks later than males, probably some of them not before mid or end of August (some juveniles probably even in mid-September). Whether they shorten the migration period and stopover times, or arrive later at the wintering grounds, or both, has to be kept open.

It is already known from ringing data from the Wand SW-European stopover sites, that the migration strategy of juveniles is quite different (summary see Le Nevé et al. 2018). After leaving the breeding grounds, they migrate in a more westerly direction to the Low Countries and France, and then along the Atlantic coast to the south. The proportion of juveniles is very high in the Low Countries and northern France, and decreases constantly towards Spain and SW-Portugal (Le Nevé et al. 2018). This matches with our geolocator data on adult males: The Loire estuary was the northernmost site with stopovers of >2.5 days, and refuelling took place mainly on the Iberian Peninsula.

Whether juvenile birds arrive later at the wintering grounds can be assumed, but would have still to be studied. The same concerns site fidelity of birds at the wintering grounds and the question, whether AW return to those wintering sites where they did spend their first winter.

## Acknowledgements

We are very grateful for the funding of the study by the NABU BirdLife Germany (for the field work) and the Förderverein Naturschutz Peenetal (for the geolocators).

We thank Ivan Bogdanovich, Maxim Koloskou, Dima and Daniil Zhuravlev from Belarus, Sven Baumung, Yves Brendahl, Filibert Heim, Benjamin Herold, Ulf Kraatz, Klaus Nigge, Andreas Prott, Cosima Tegetmeyer, Florens Waldschmidt and Landelin Winter from Germany, Toms Endziņš, Aivis Gulbis, Roberts Jansons, Valts Jaunzemis and Elza Zacmane from Latvia, Jurate Zarankaite from Lithuania and Grzegorz Kiljan from Poland for participating in the fieldwork.

We are grateful to Alexandre Vintchevski (former APB – BirdLife Belarus) for his very important organisational support and Maxim Nemtchinov (APB) for the excellent preparation and organisation of fieldwork in the Servech mire. Zymantas Morkvenas and Rita Griniene (Baltic Environmental Forum BEF) organized our stay in Lithuania and enabled very valuable insights into the AW translocation operation at Zuvintas biosphere reserve within the EU-LIFE project MagniDucatusAcrola.

David Miguélez (Fundacion Global Nature) gave very valuable advice for the identification of potential stopover sites (“informed guesses”) in central Spain. Jens Hering gave valuable advice on habitats and potential stopover sites in Algeria.

We thank James Fox (Migrate Technology Ltd.) not only for providing us with excellent geolocators without any malfunctions, but also for his very quick and friendly, great support in offloading the data from the geolocators. We are also grateful to Janne Ouwehand (University of Groningen), who contributed very helpful information regarding harness material and harness construction for the geolocators.

---

**Annex 3:**
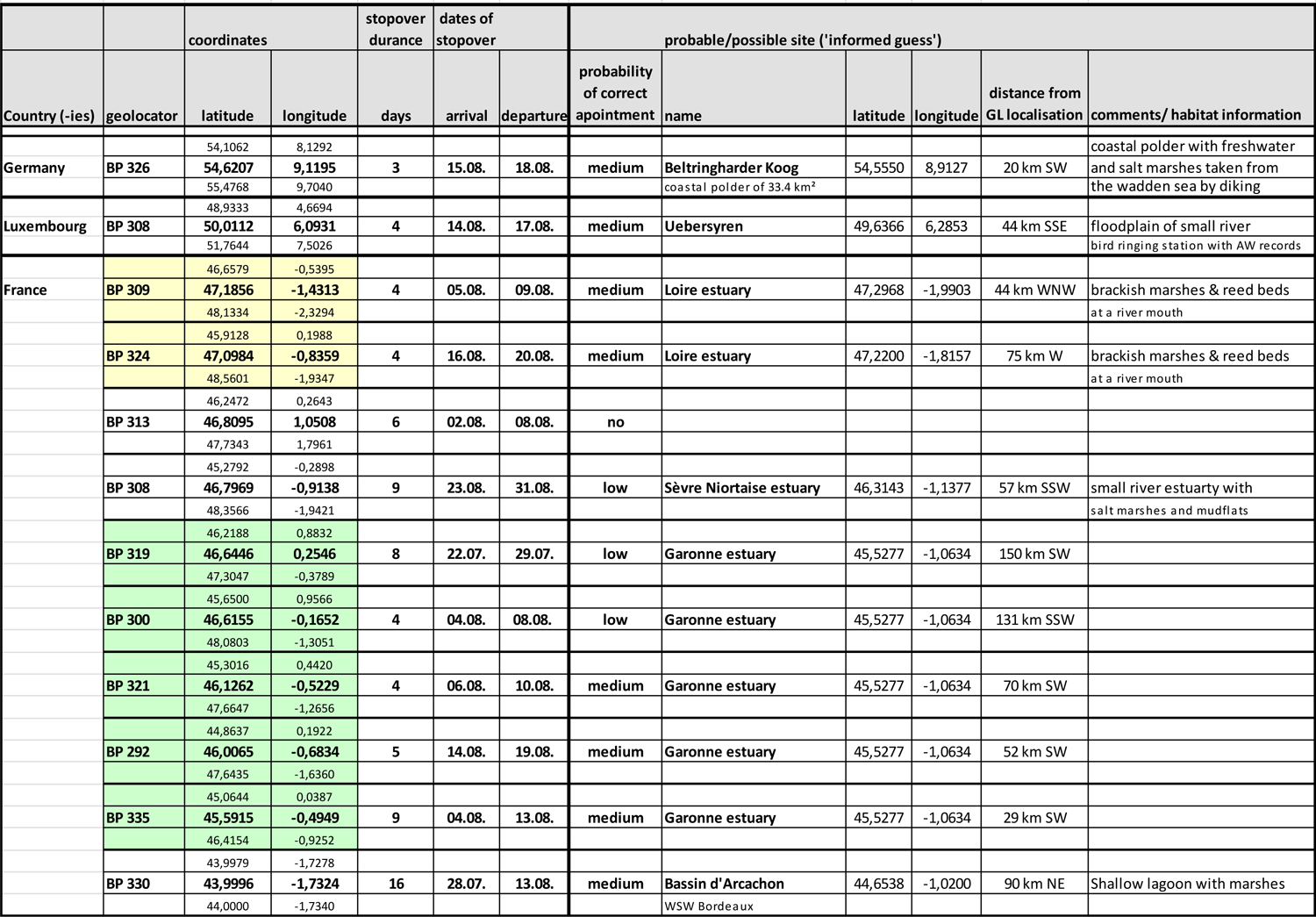
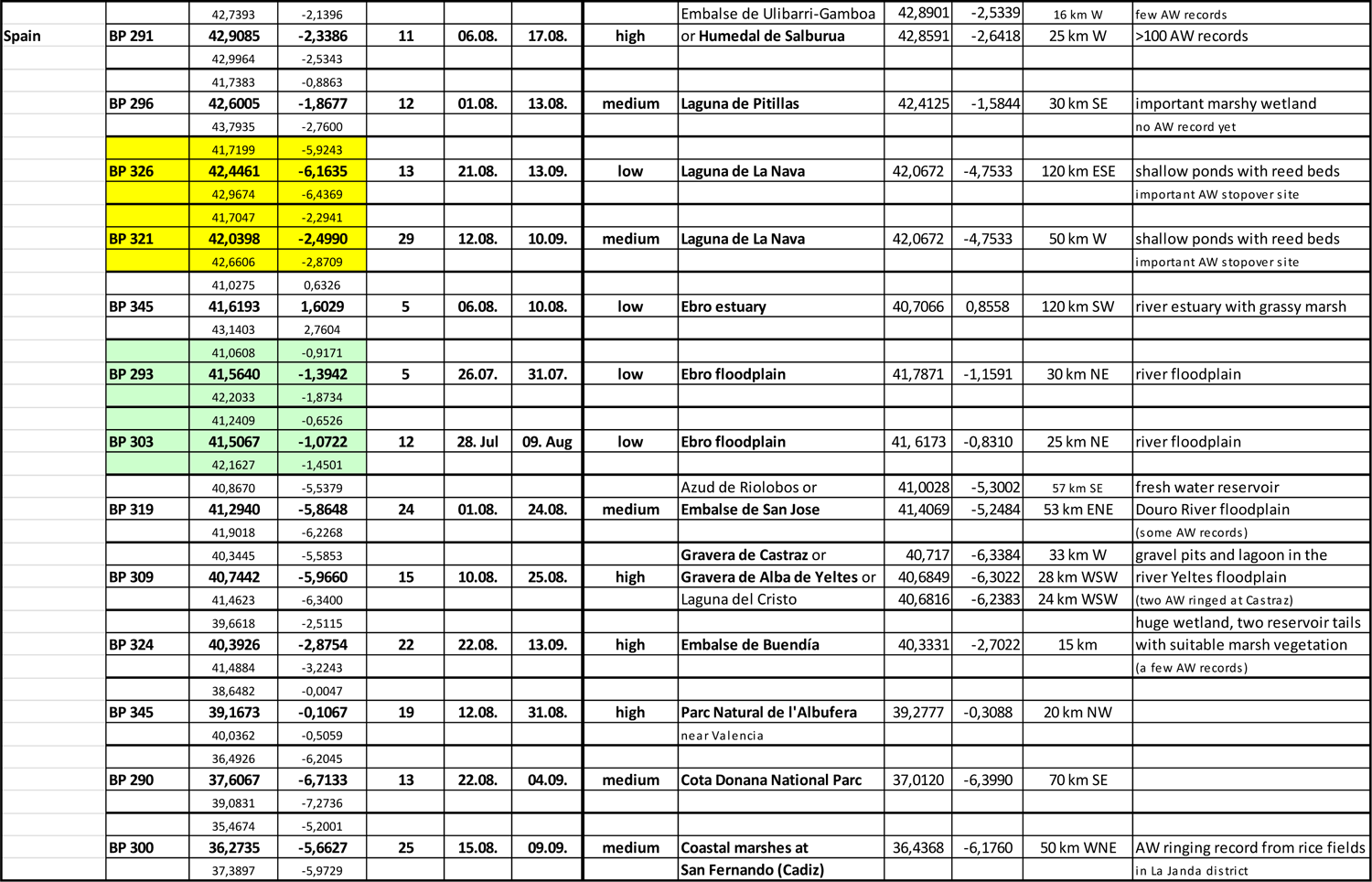
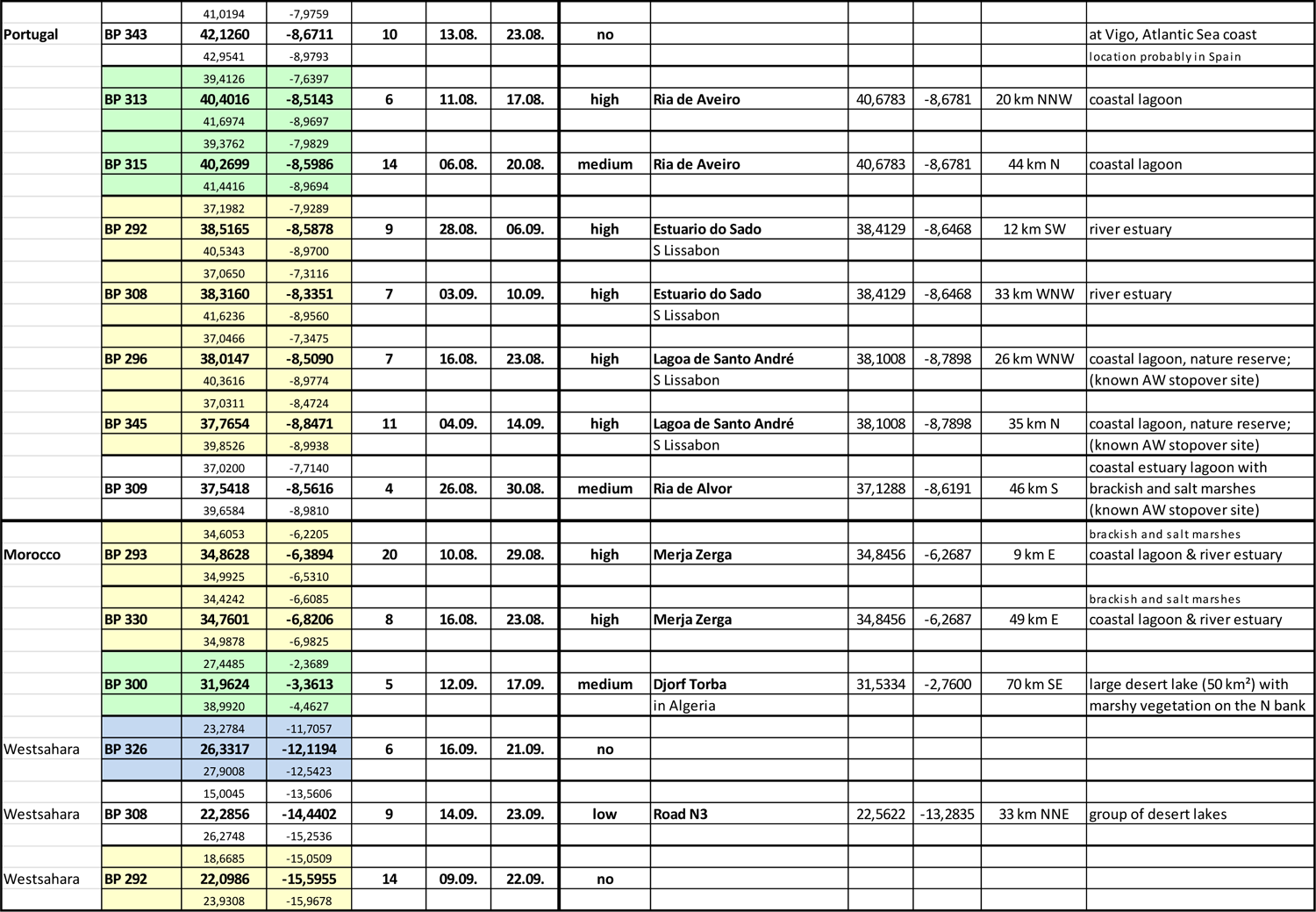
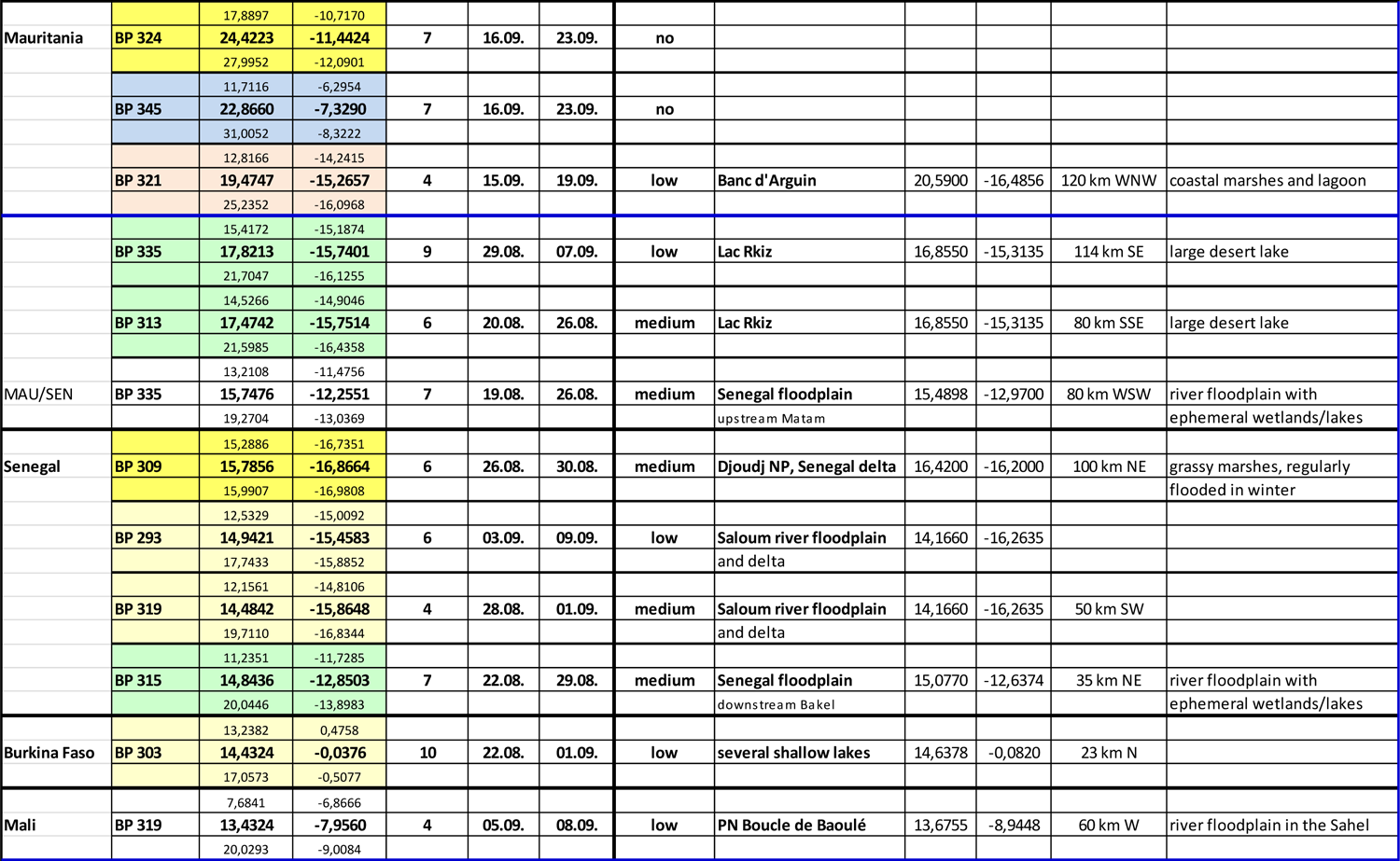
AW stopovers on autumn migration (>2.5 days)

**Annex 4:**
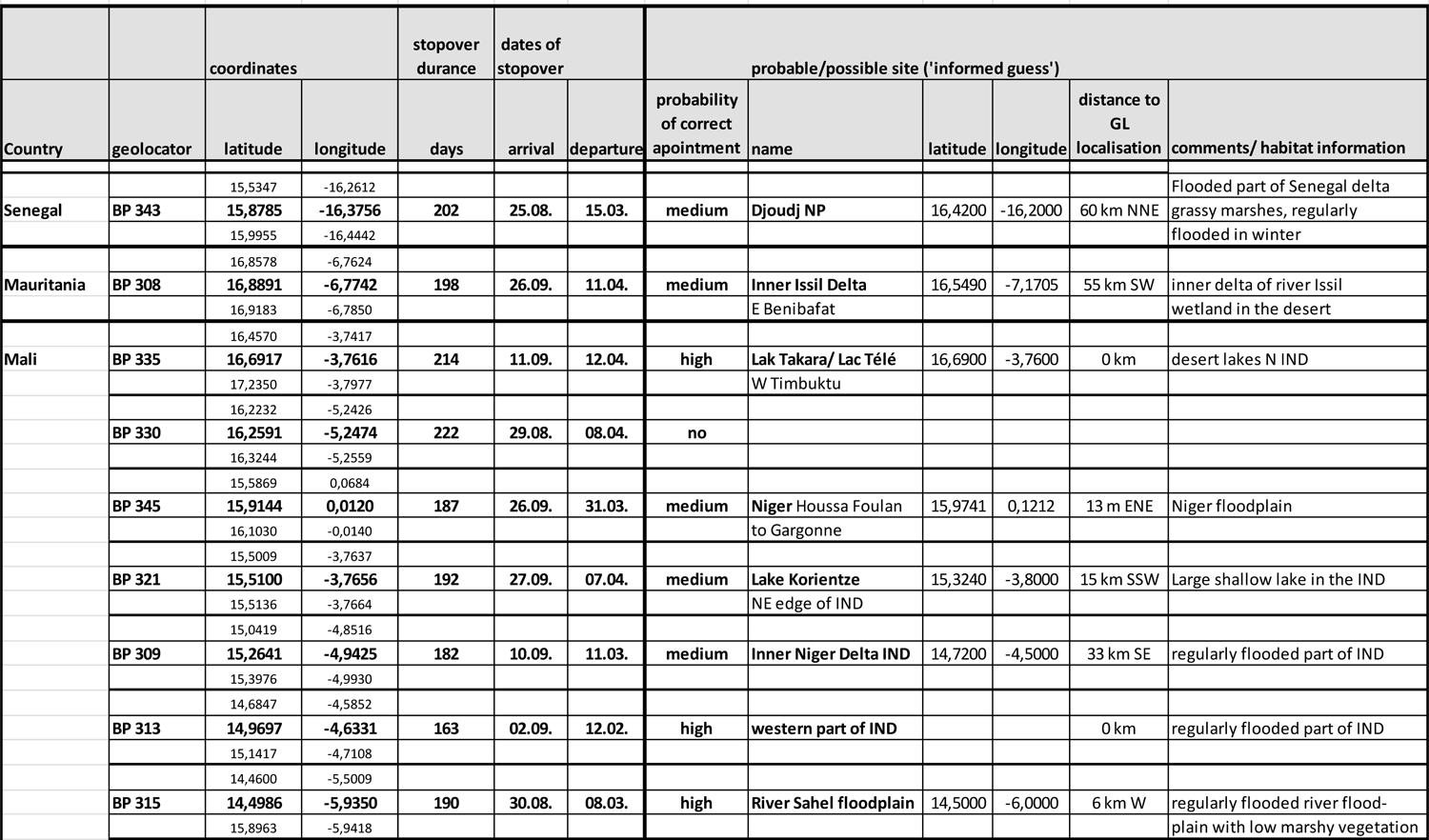
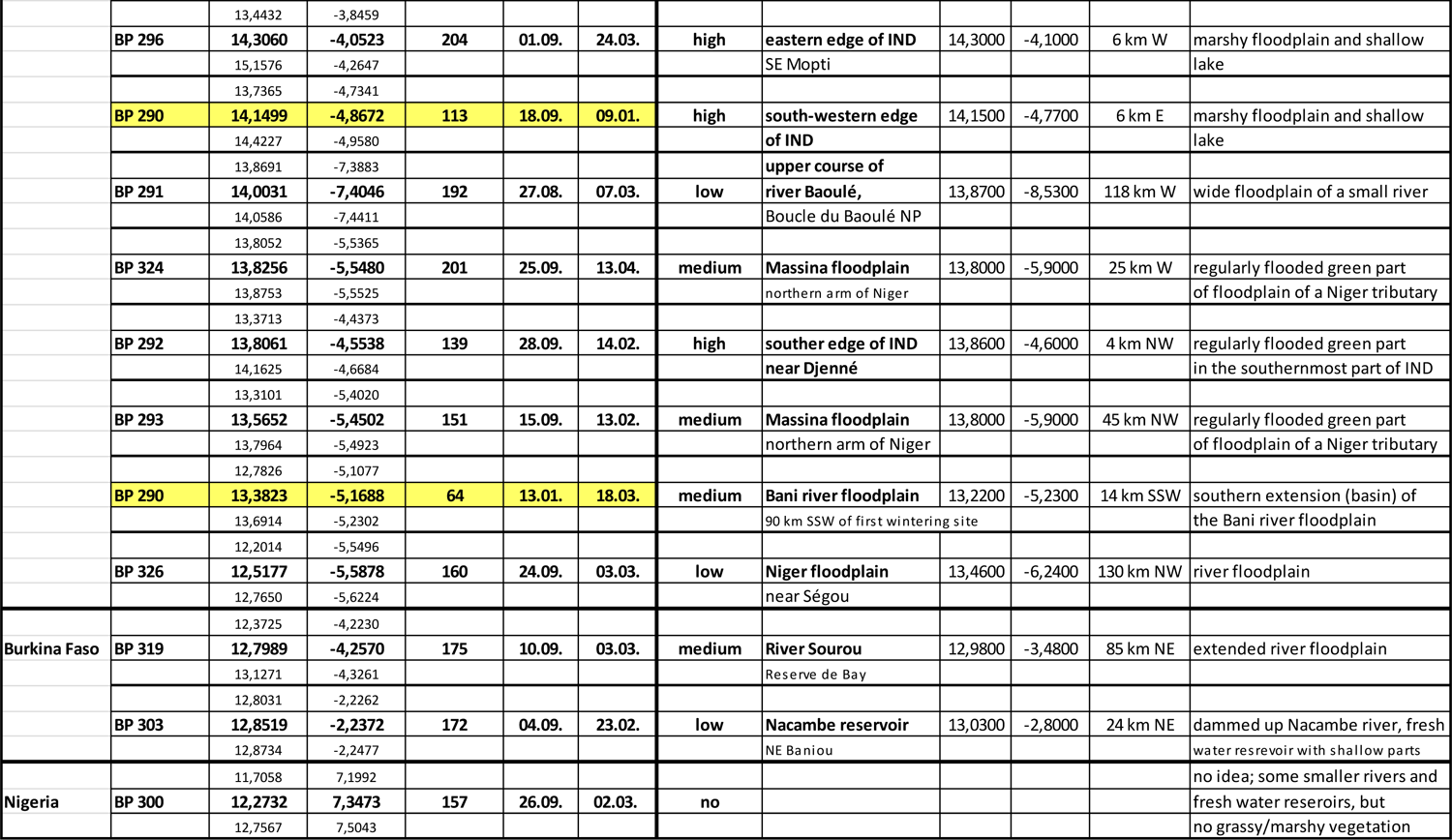
AW wintering sites (>60 days October-February)

**Annex 5:**
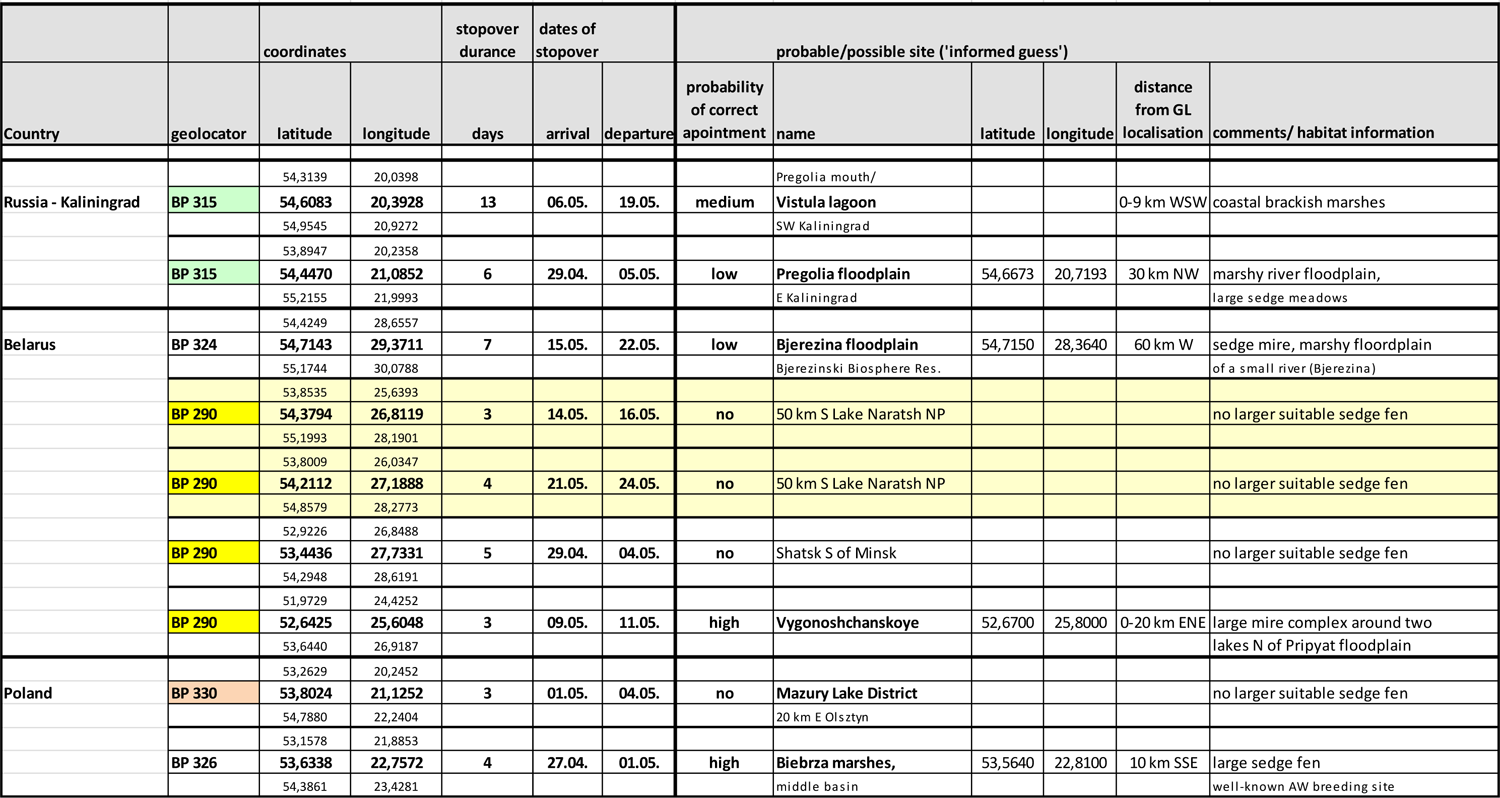
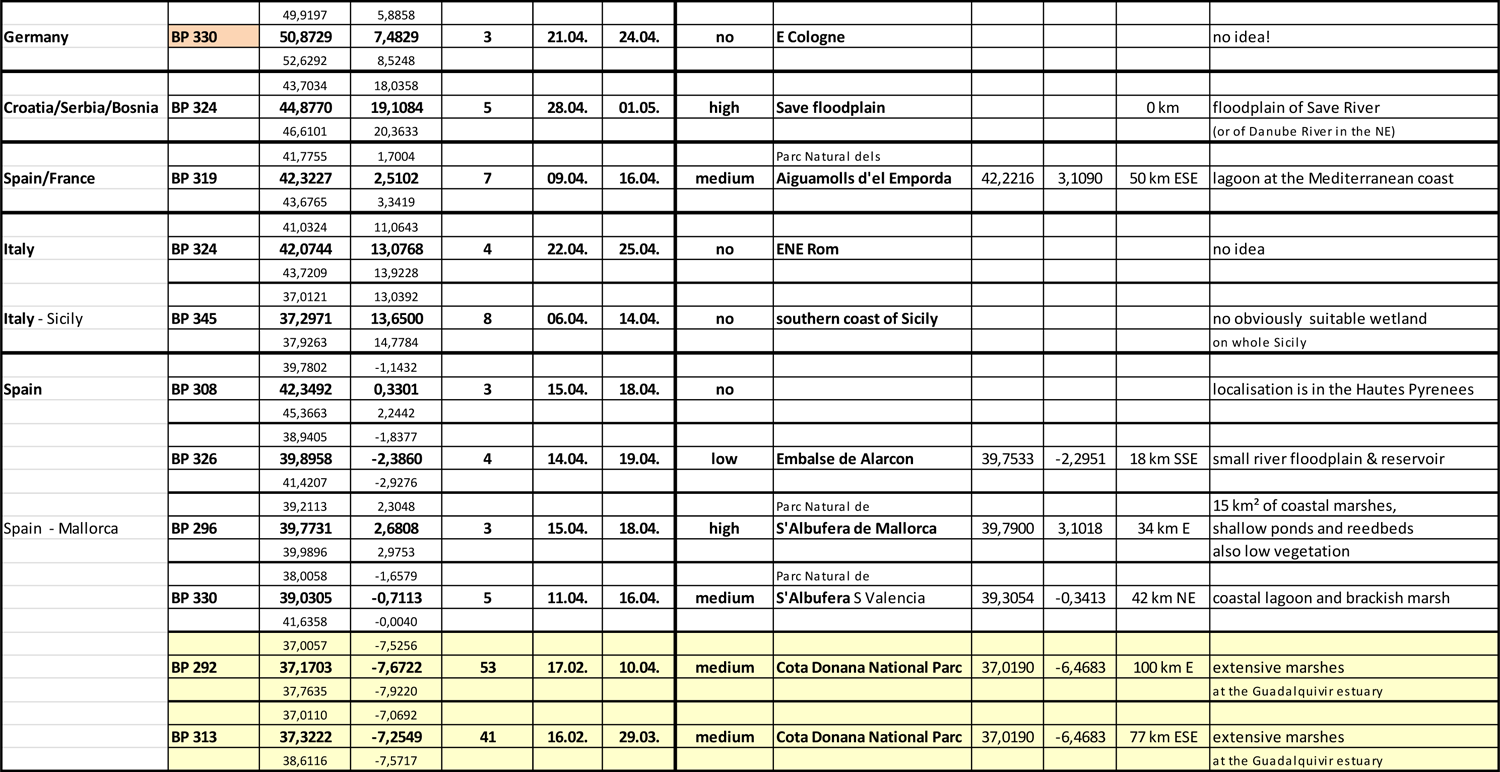
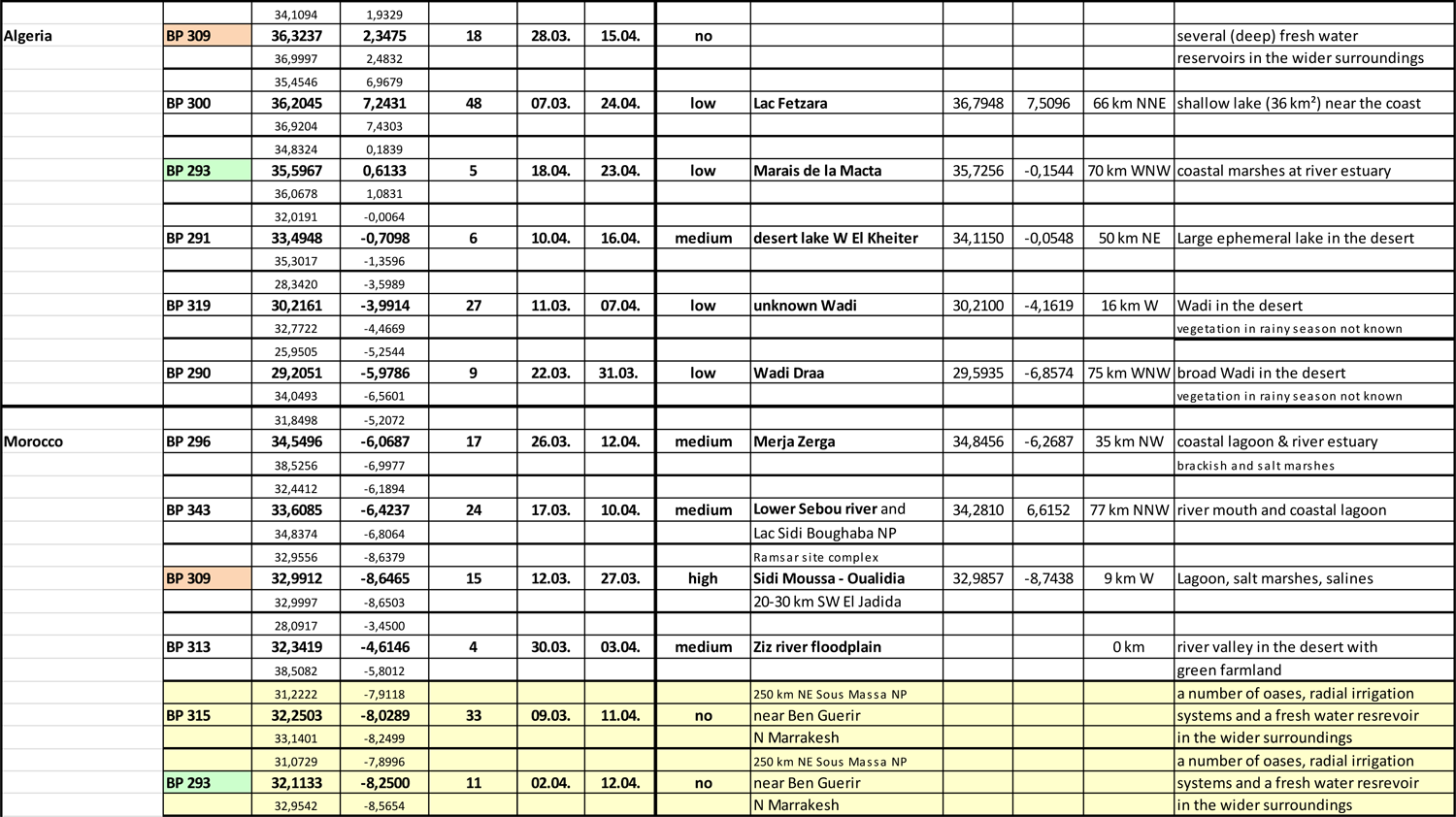
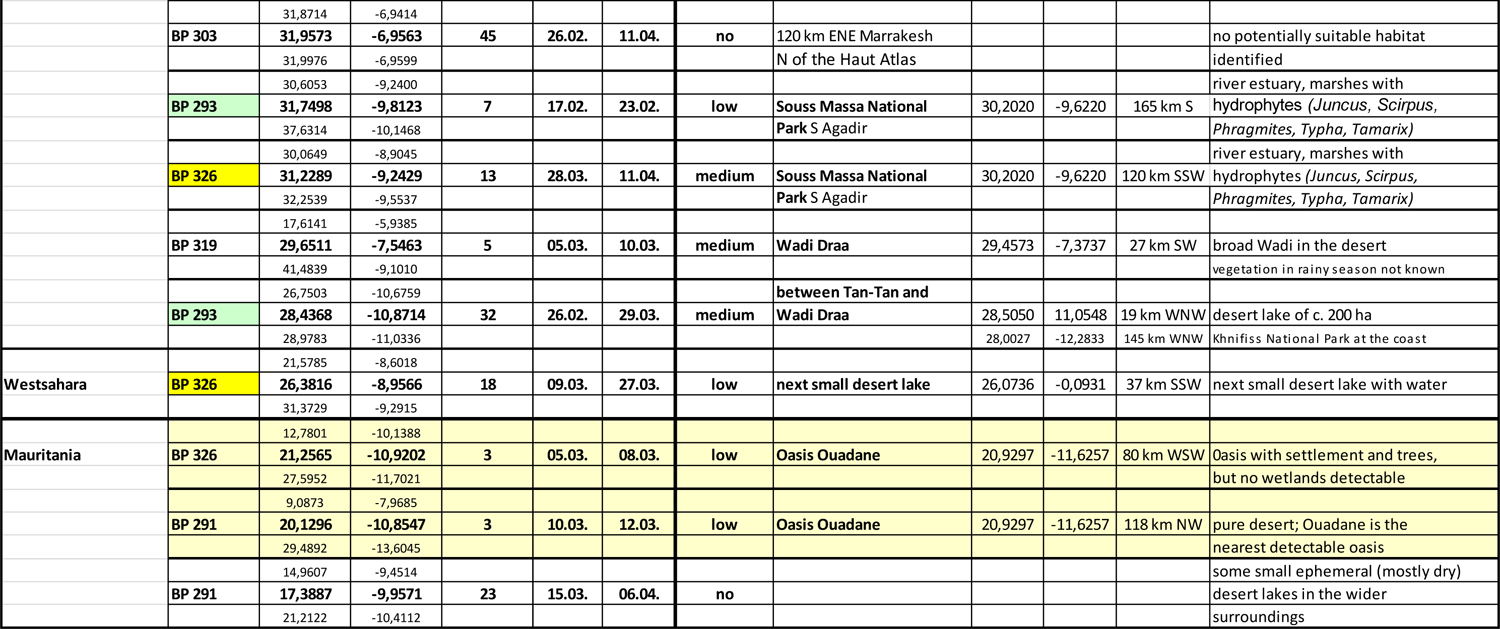
AW stopover sites on spring migration (>2.5 days)

**Annex 6:**
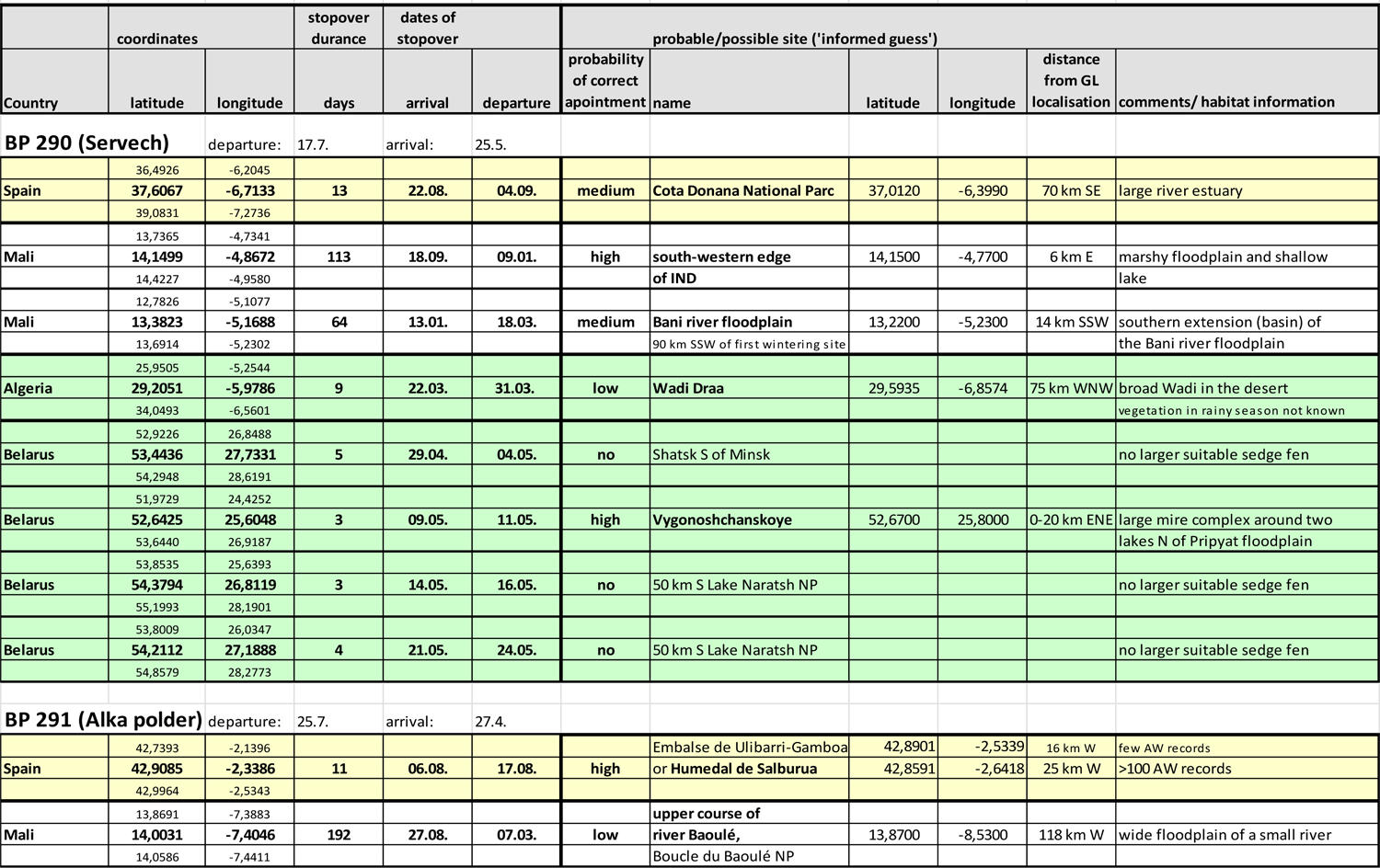
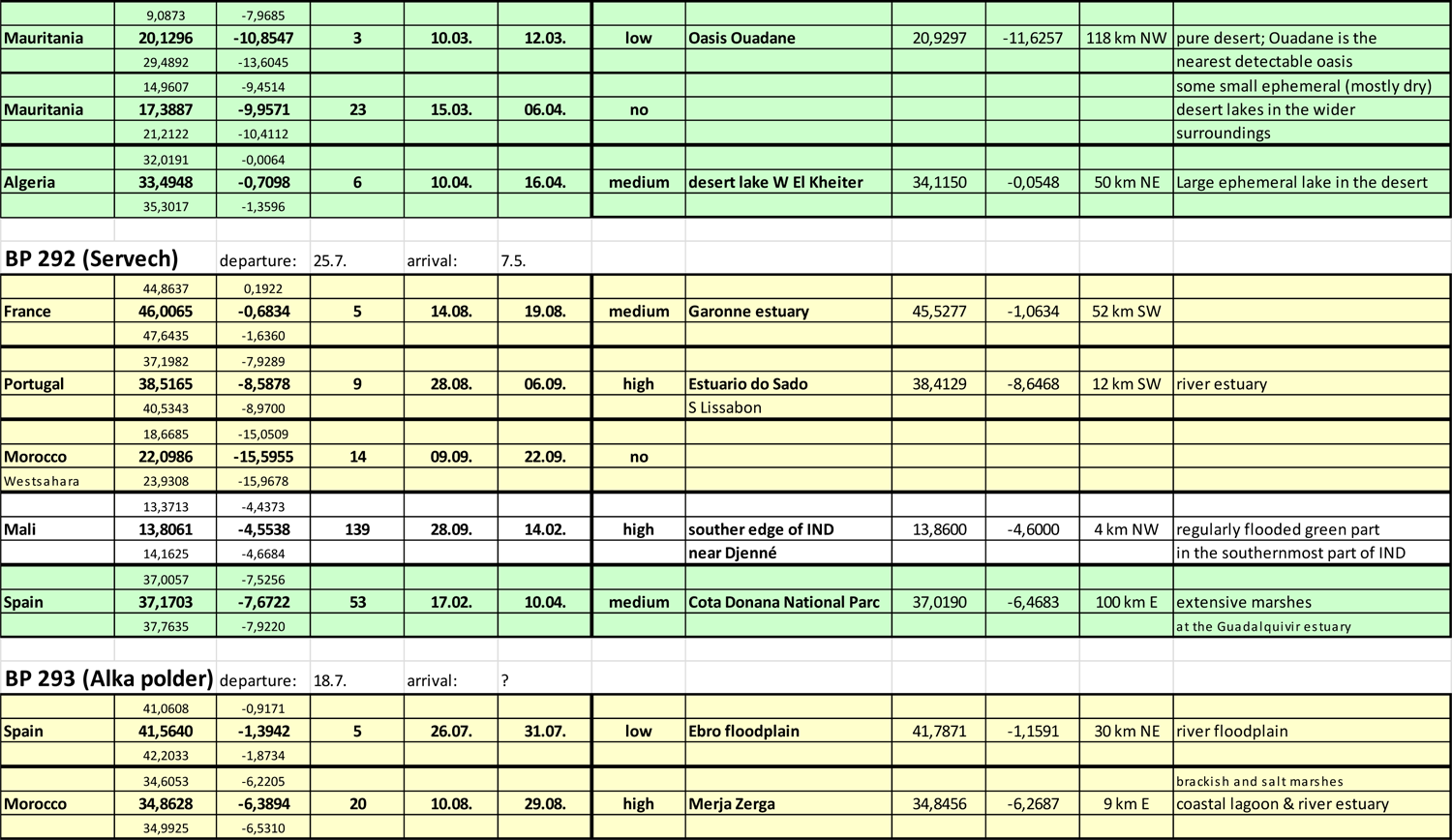
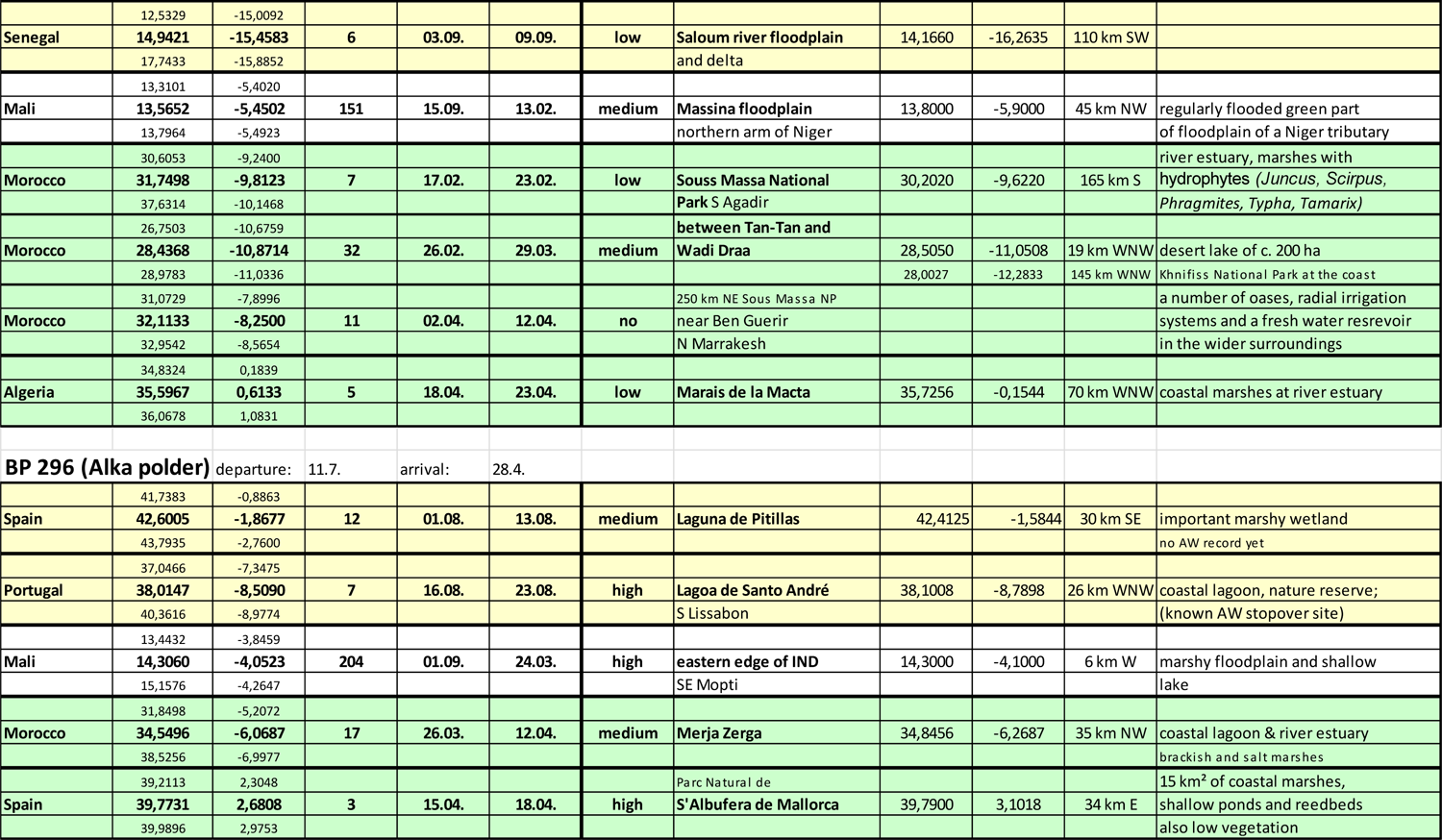
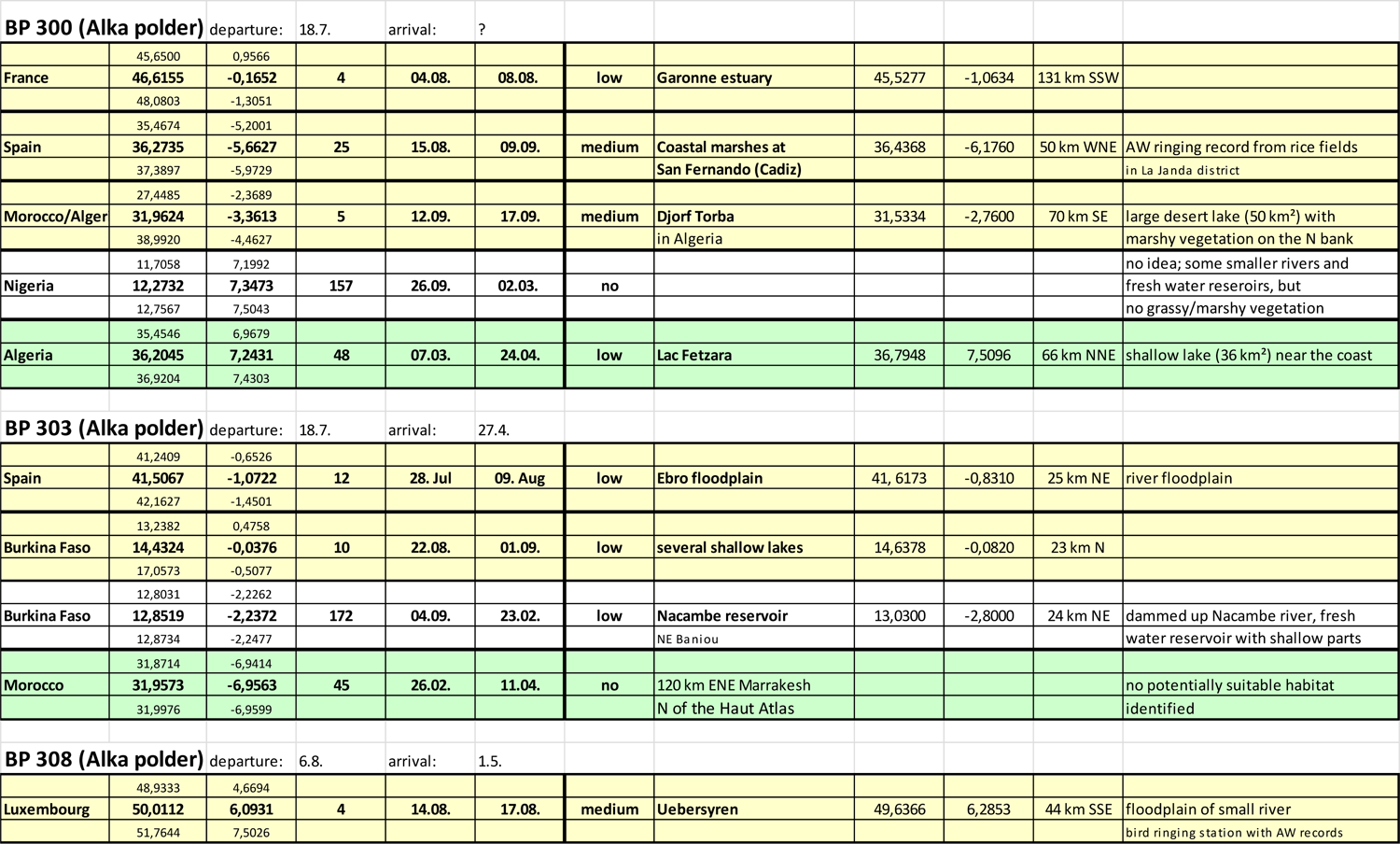
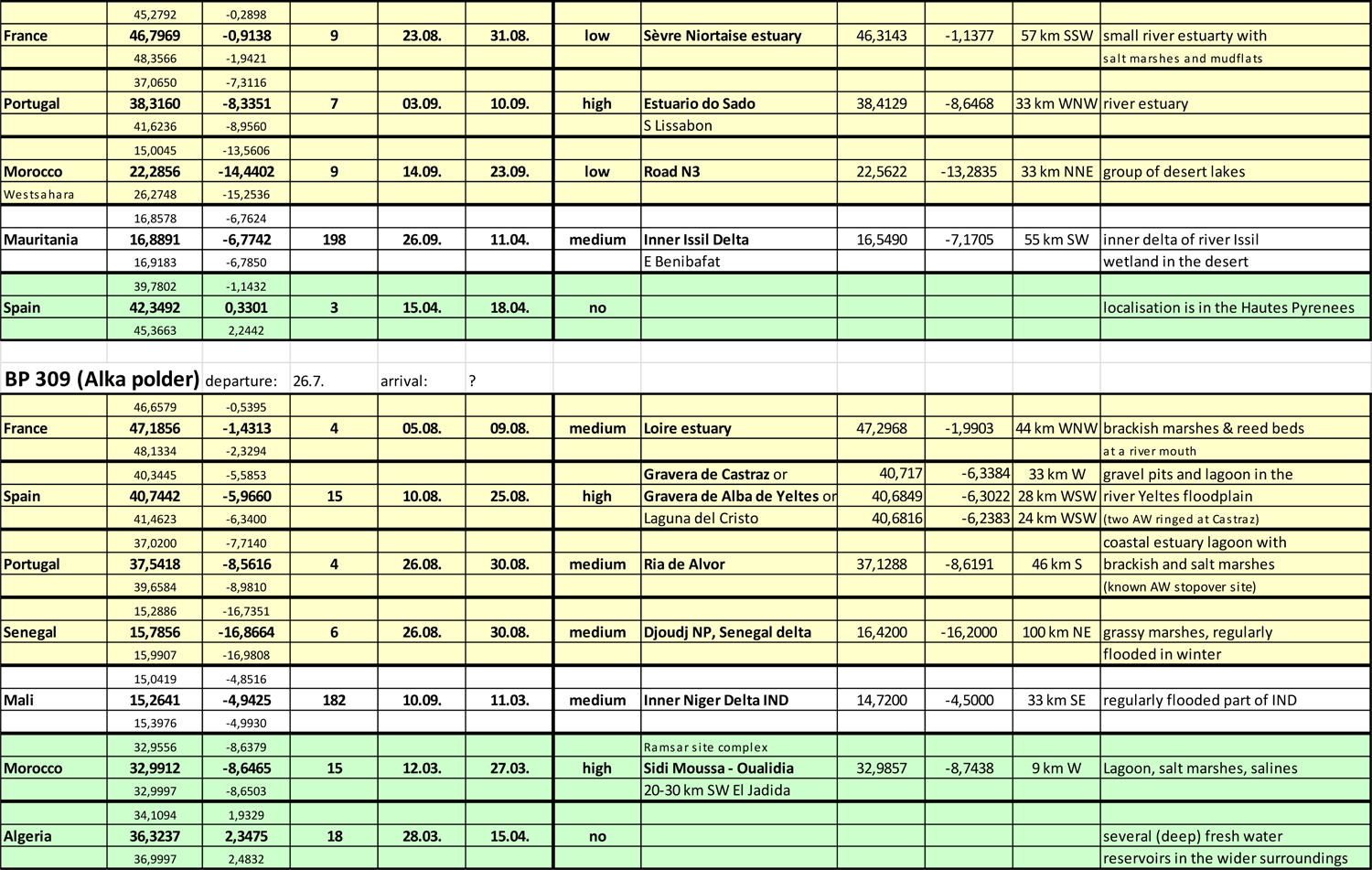
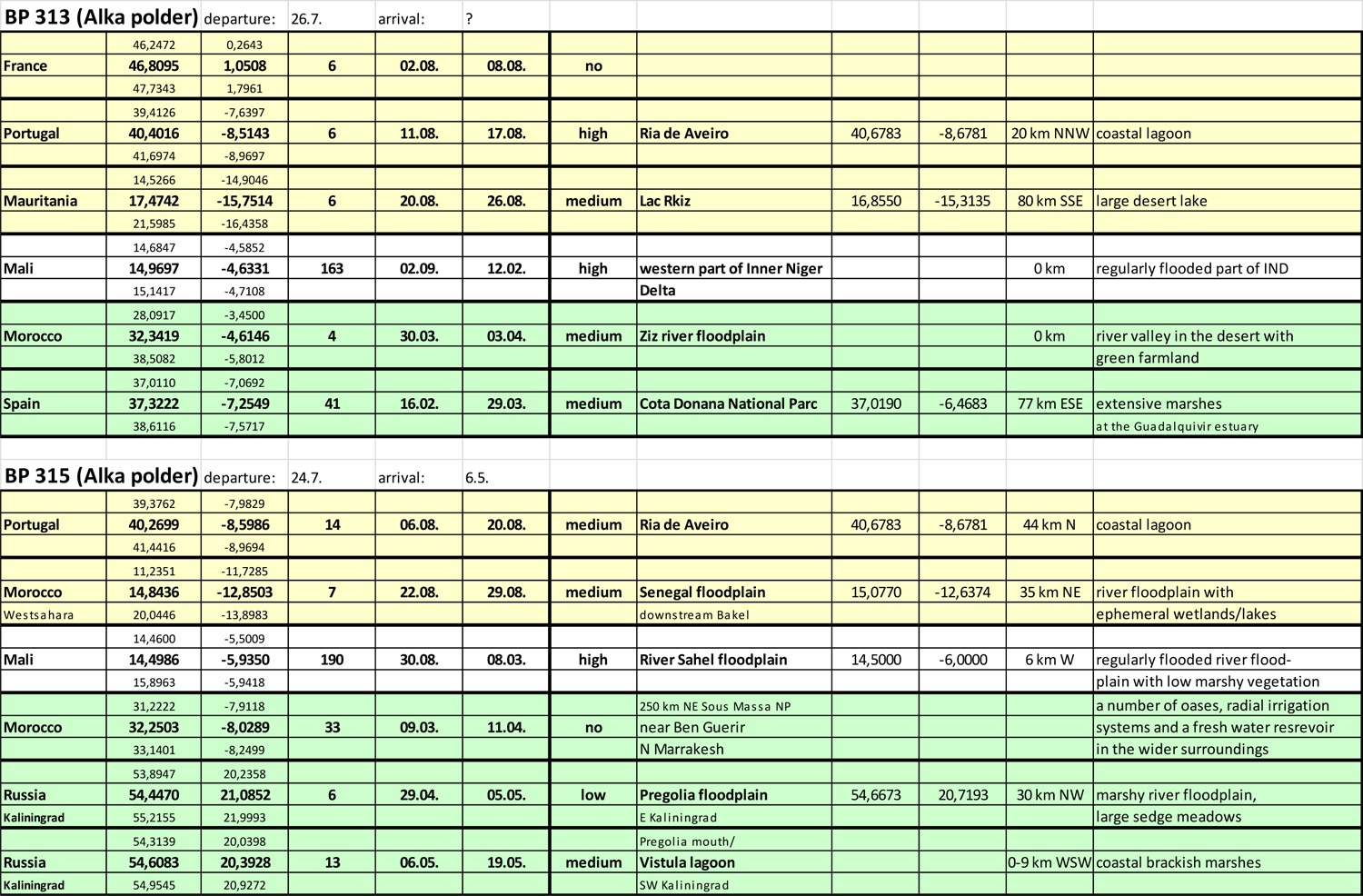
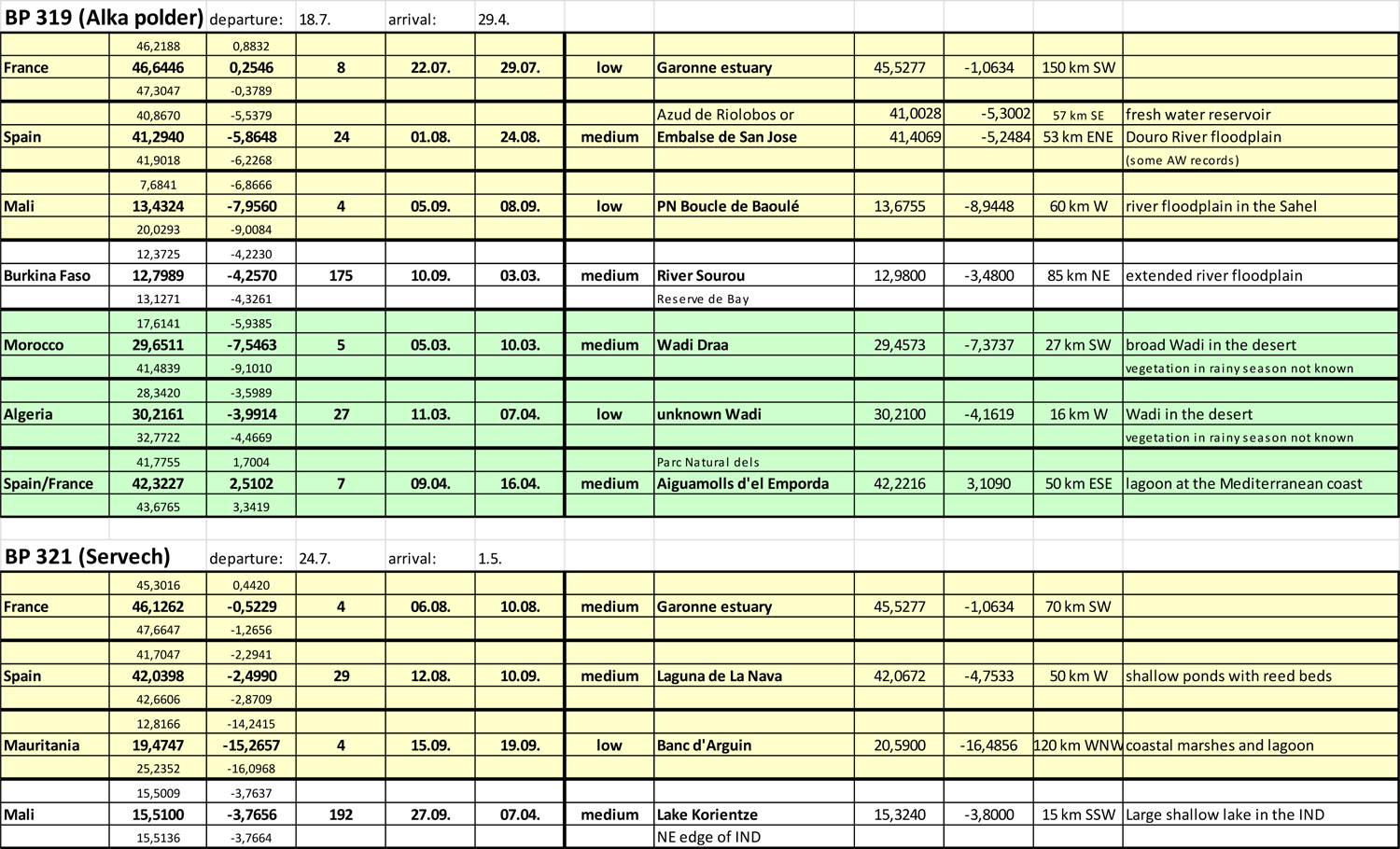
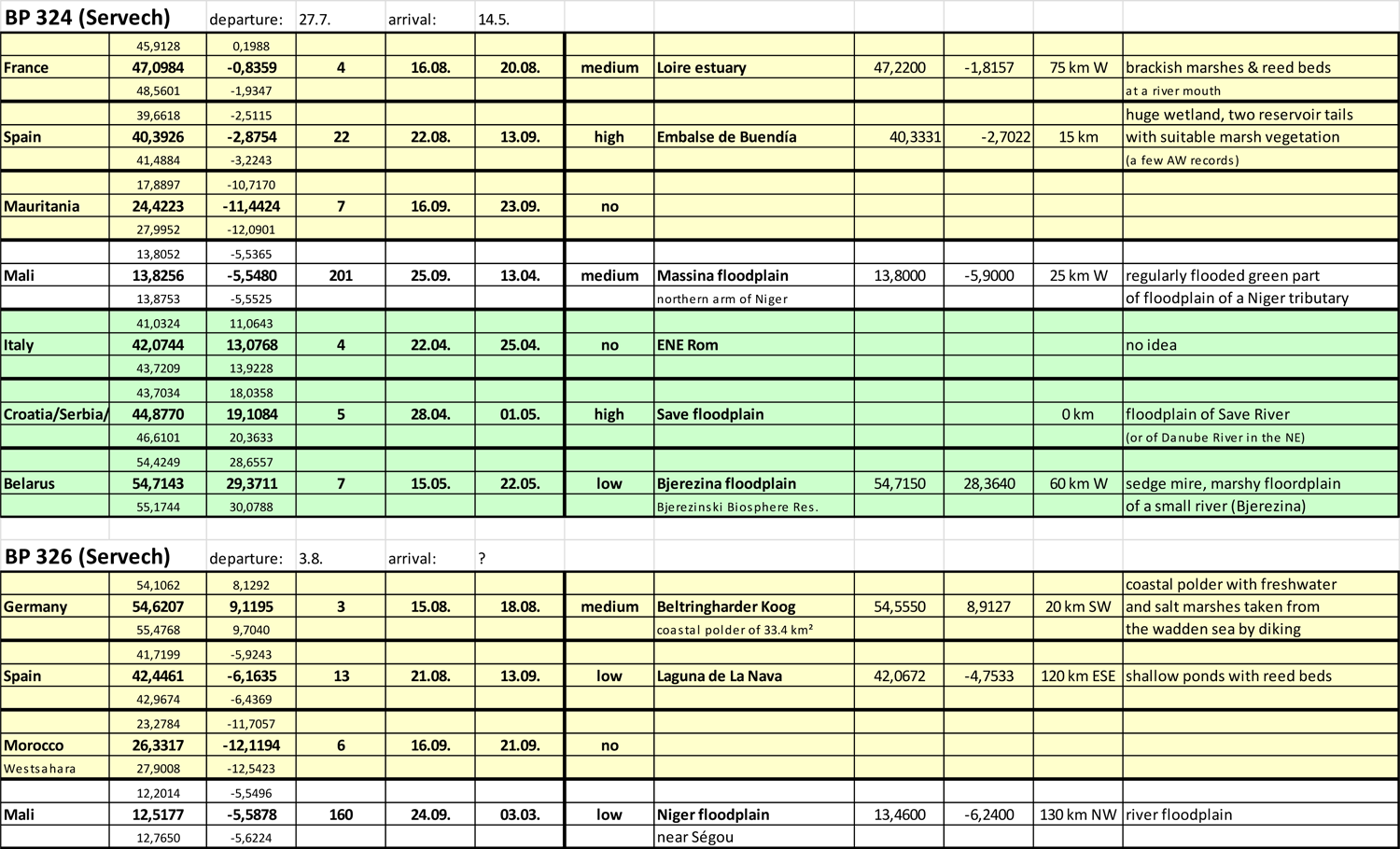
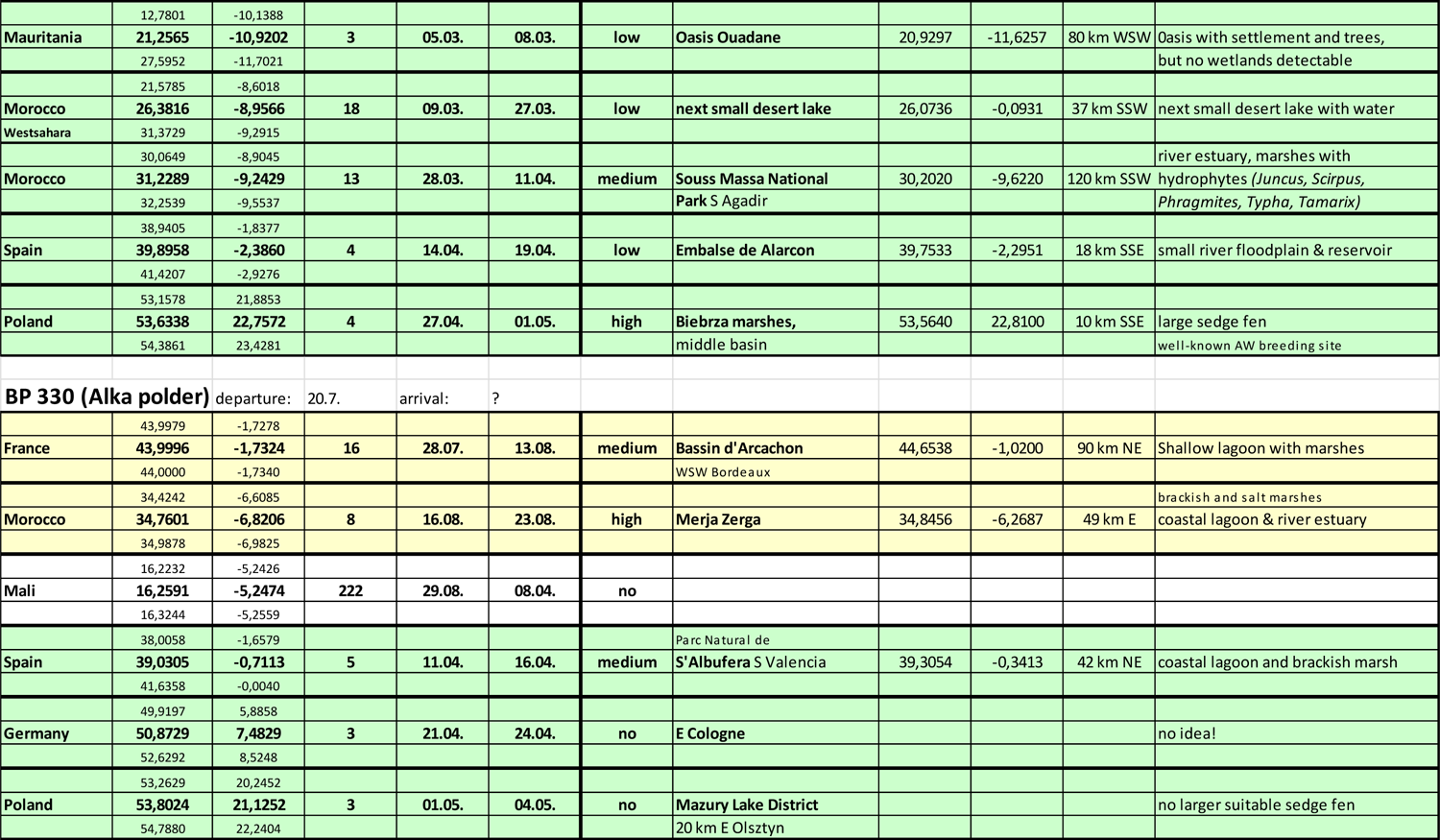
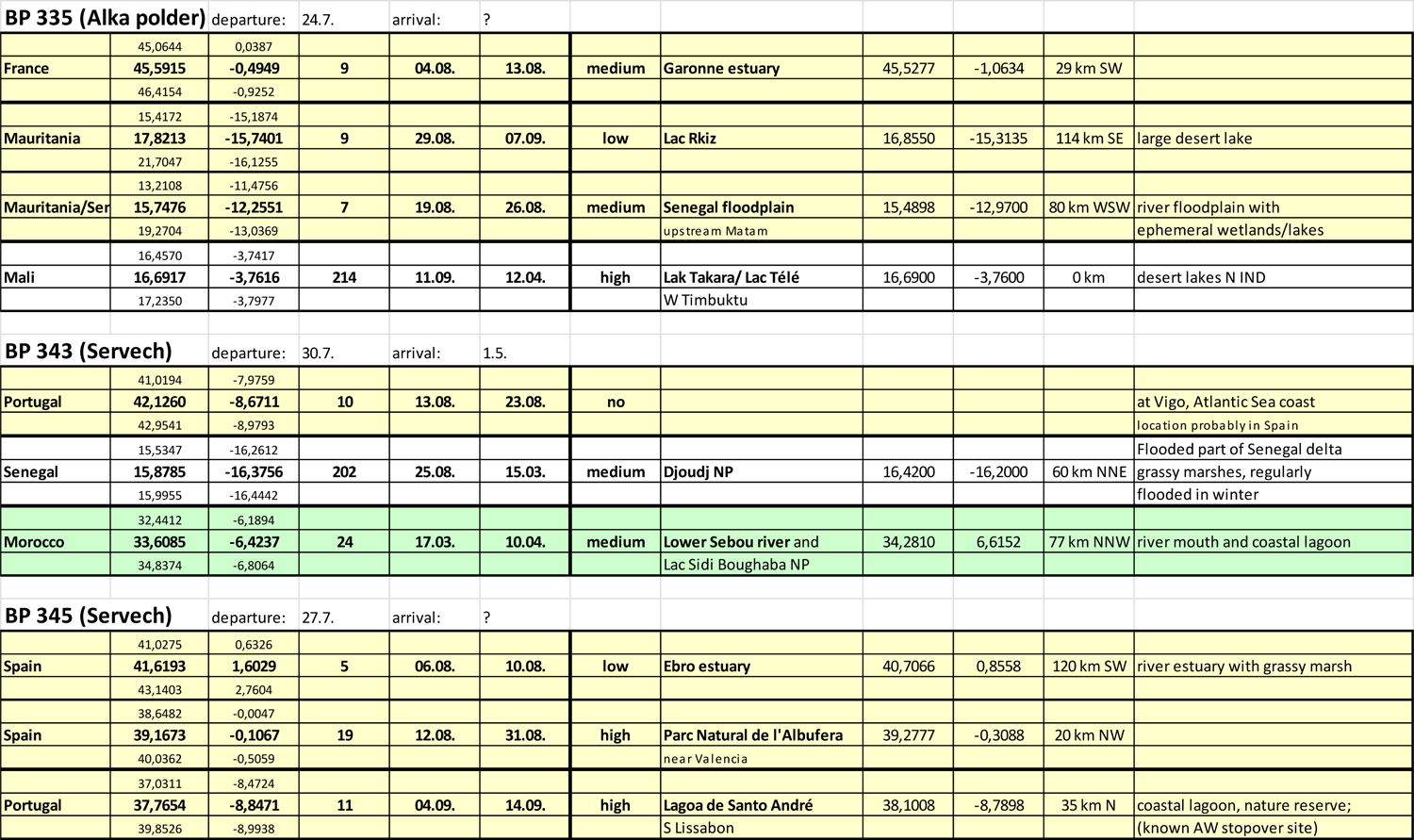
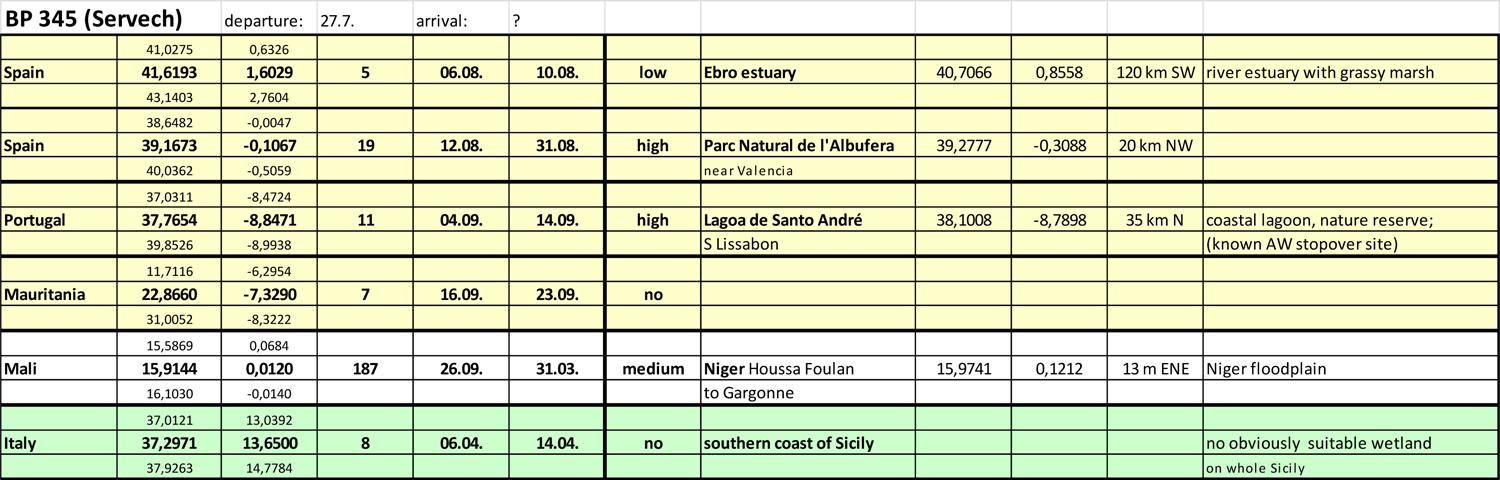
Stopover and wintering sites according to geolocator numbers.

